# Cleavage of the pseudoprotease iRhom2 by the signal peptidase complex reveals an ER-to-nucleus signalling pathway

**DOI:** 10.1101/2022.11.28.518246

**Authors:** Iqbal Dulloo, Michael Tellier, Clémence Levet, Anissa Chikh, Boyan Zhang, Catherine M Webb, David P Kelsell, Matthew Freeman

## Abstract

iRhoms are pseudoprotease members of the rhomboid-like superfamily and are cardinal regulators of inflammatory and growth factor signalling; they function primarily by recognising transmembrane domains of their clients. Here we report an unexpected, and mechanistically distinct, nuclear function of iRhoms. iRhom2 is a non-canonical substrate of the signal peptidase complex (SPC), the protease that removes signal peptides from secreted proteins. Cleavage of iRhom2 generates an N-terminal fragment that enters the nucleus and modifies the cellular transcriptome. We observed elevated nuclear iRhom2 in skin biopsies of patients with psoriasis and tylosis with oesophageal cancer (TOC); increased SPC-mediated iRhom2 cleavage in a psoriasis model; and overlapping transcriptional signatures between psoriasis and expression of the iRhom2 N-terminus. This work highlights the pathophysiological significance of this SPC-dependent ER-to-nucleus signalling pathway, and is the first example of a rhomboid-like protein that mediates protease-regulated nuclear signalling.

## INTRODUCTION

The rhomboid-like superfamily of membrane proteins comprises the originally-discovered rhomboid intramembrane serine proteases, and multiple pseudoproteases which, despite being widely conserved, have lost their protease activity (Freeman, 2014; Lemberg and Adrain, 2016). Pseudoenzymes were once assumed to be functionally dead evolutionary remnants, but they are emerging as an important class of proteins with significant biological functions (Adrain and Freeman, 2012), and this is consistent with what is known of the diverse functions of pseudoprotease members of the rhomboid-like superfamily, the best characterised of which are iRhom1 and iRhom2 (Dulloo et al., 2019). They are now most famous as regulatory cofactors of ADAM17, a cell surface metalloprotease responsible for the release of important intercellular signalling proteins (Zunke and Rose-John, 2017). As such, iRhoms control both inflammatory signalling by the cytokine TNF, and growth factor signalling by members of the EGF family (Adrain et al., 2012; McIlwain et al., 2012; Zettl et al., 2011). iRhoms are, however, multifunctional and they also participate in, for example, the response to chronic ER stress (Dulloo et al., 2022) and viral infection (Luo et al., 2016).

Rhomboid-like proteins have a modular structure (Dulloo et al., 2019). The mechanistic theme believed to underlie all their functions is the specific recognition of, and interaction with, transmembrane domains (TMDs) of substrates and client proteins. This TMD recognition function is mediated by their conserved transmembrane core. All rhomboid-like proteins also have cytoplasmic and luminal/extracellular domains, which are less well conserved, and are presumed to mediate functions more specific to particular members of the superfamily. In the case of the iRhoms, these include a long cytoplasmic N-terminus, and a large luminal loop between TMD1 and TMD2, called the iRhom homology domain (IRHD) (Lemberg and Freeman, 2007). The cytoplasmic N-terminus is predicted to be largely unstructured but has important regulatory properties, interacts with several accessory factors and signalling proteins, as well as being the site of post-translational modifications (Grieve et al., 2017; Kunzel et al., 2018; Oikonomidi et al., 2018) and disease-associated mutations (Blaydon et al., 2012).

We and others have previously noted that C-terminally tagged iRhoms exist in two forms: the full-length protein, and a shorter fragment (Adrain et al., 2012; Christova et al., 2013; Grieve et al., 2017; Kunzel et al., 2018; McIlwain et al., 2012; Nakagawa et al., 2005; Zettl et al., 2011) whose size suggests that the N-terminal cytoplasmic domain is deleted. Indeed, the N-terminal domain may not be essential for some iRhom functions: its deletion is reported not to abolish function but instead to cause elevated constitutive ADAM17 activity in mammalian cells (Hosur et al., 2014; Maney et al., 2015) and, in *Drosophila,* a wing phenotype that suggests hyperactivity (Nakagawa et al., 2005). Nevertheless, whether shorter forms of iRhoms have any physiological function remains unexplored.

Pursuing this question, we have discovered that endogenous iRhom2 undergoes partial proteolytic cleavage to generate three stable forms: the full-length protein, and both N- and C-terminal fragments. We have identified the protease responsible for this iRhom2 cleavage as the signal peptidase complex (SPC). SPC is primarily the protease responsible for the removal of signal peptides from proteins entering the ER (Jackson and Blobel, 1977; Liaci et al., 2021), but in this case cleaves iRhom2 non-canonically, adjacent to its first transmembrane domain (TMD). The consequence of SPC cleavage is the release of the N-terminal domain of iRhom2 and its translocation to the nucleus, where it modifies the cellular transcriptome. Strikingly, the pathophysiological significance of this newly uncovered ER-to-nucleus pathway is indicated by our observation that skin biopsies particularly from patients with psoriasis or the genetic syndrome tylosis with oesophageal cancer (TOC) show elevated nuclear iRhom2. There is also significant overlap between genes differentially expressed in lesional psoriatic skin and those upregulated by nuclear iRhom2, accompanied by higher expression levels of iRhom2 and several components of the SPC. Overall, this work expands our understanding of the spectrum of functions of both rhomboid-like proteins and SPC.

## RESULTS

### A cleaved N-terminal fragment of iRhom2 translocates to the nucleus

A HEK293T cell line in which endogenous iRhom2 was C-terminally tagged with a 3XHA tag by CRISPR knock-in confirmed previous observations (Kunzel et al., 2018) that endogenous iRhom2 exists as both full-length and a shorter C-terminal forms. The identity of both bands as iRhom2 was validated with two different siRNAs against iRhom2 (Fig. 1a). Similarly, overexpression of C-terminally 3XHA-tagged iRhom1 and iRhom2 led to the generation of both full-length protein (± 100 KDa; iR1/2-FL), and a shorter C-terminal form of approximately 50 kDa (iR1/2-CT) (Fig. 1b, top panel). Using an antibody specific to the N-terminal cytoplasmic domain (Fig. 1b, bottom panel), we detected a N-terminal form of iRhom2 of approximately 45 kDa (iR2-NT). The endogenous existence of this N-terminal fragment was confirmed in wild-type but not iRhom2 knockout mouse lung tissue (Supplementary Fig. 1a). We conclude that iRhoms exist in three major forms – the full-length protein, and N- and C-terminal fragments, whose combined sizes suggest that they are products of proteolytic cleavage.

**Fig. 1.**
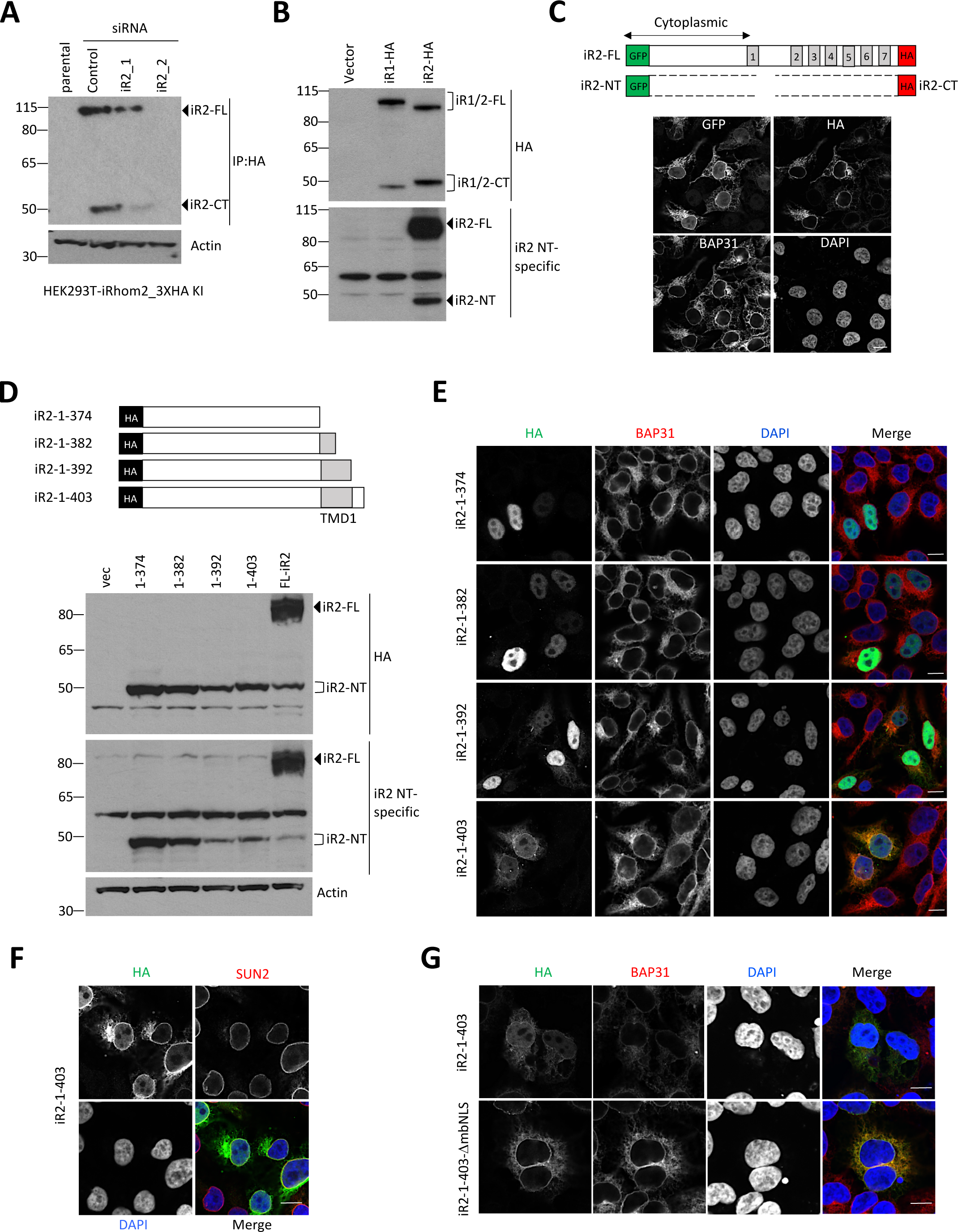
A cleaved iRhom2 fragment translocates to the nucleus. **a** Endogenously tagged iRhom2-HA was detected by first immunoprecipitation (IP: HA), followed by immunoblotting with HA antibody in lysates from HEK293T cells with 3XHA tag inserted in frame at the C-terminus of iRhom2. Cells were transfected with control siRNA or two different siRNAs against iRhom2 for 72 h before harvest. **b** Levels of iRhom1 and iRhom2 were analysed by immunoblotting in lysates from HEK293T cells transiently transfected for 36 h with iRhom1-3XHA and iRhom2-3XHA, using either anti-HA or an iRhom2 N-terminal-specific antibody. **c** Schematic showing expected products from GFP-iRhom2-3XHA (top). Immunofluorescence of GFP-iRhom2-3XHA transfected in HEK293T cells for 36hrs. Cells were stained for GFP, HA, BAP31 as an ER marker and DAPI to show the nucleus. Scale bar =10 *μ*m. **d** Schematic showing truncated iRhom2 mutants with N-terminal 2XHA tag (top). The size of the N-terminal fragment from full-length iRhom2 was compared with iRhom2 truncations by immunoblotting in lysates from HEK293T cells transiently transfected for 36 h using either anti-HA or iRhom2 N-terminal specific antibodies. **e** Immunofluorescence of truncated iRhom2 mutants transfected in HEK293T cells for 36hrs. Cells were stained for HA (green), BAP31 to label ER (red) and DAPI to label nuclei (blue). Scale bar =10 *μ*m. **f-g** Immunofluorescence of iR2-1-403 (**f-g**) and iR2-1-403_ΔmbNLS (**g**) transfected in HEK293T cells for 36hrs. Cells were stained for HA (green), either SUN2 as a nuclear membrane marker (red) (**f**), or BAP31 (red) (**g**) and DAPI (blue). Scale bar =10 *μ*m. Immunofluorescence and immunoblots data are representative of 2-3 independent experiments.

Treatment of cells overexpressing iRhom2 with the proteasomal inhibitor MG-132 (Supplementary Fig. 1b) or the lysosomal inhibitors chloroquine and 3-MA (Supplementary Fig. 1c) did not prevent the generation of the shorter C-terminal iRhom2 fragment, suggesting that the fragments were not products of protein degradation. We also asked whether the N- and C-terminal forms of iRhom2 were stable in cells. Cycloheximide chase experiments showed that the cleaved iRhom2 protein had a half-life >4hrs, while the full-length protein half-life was approximately 2hrs (Supplementary Fig. 1d-e). These data confirm that iRhoms proteins exist in multiple stable forms, apparently due to the proteolytic cleavage of the full-length protein into an N-terminal and a C-terminal fragment.

Full-length iRhom2 is a polytopic membrane protein with two known major cellular locations: the ER and the plasma membrane (Dulloo et al., 2019; Kunzel et al., 2018; Oikonomidi et al., 2018). N-linked glycans attached to ER proteins are sensitive to the deglycosidase Endo H, while the glycans on post-Golgi proteins are insensitive to Endo H but can be removed by PNGase F. In cells expressing C-terminally tagged iRhoms, Endo H treatment caused a downshift of both the full-length and the C-terminal iRhom2 fragment, indicating that the C-terminal iRhom2 fragment is predominantly localised in the ER (Supplementary Fig. 1f). Consistent with this, in immunofluorescent staining, the full-length (GFP- and HA tagged) and C-terminal iRhom2 fragment (HA tagged) colocalised with the ER resident protein BAP31, clearly visible in the ER and the nuclear envelope (which is contiguous with the ER) (Fig. 1c). To our surprise, however, the N-terminal fragment (iR2-NT), labelled with the GFP tag, also showed weak but reproducible diffuse staining in the nucleus (Fig. 1c). A different iRhom2 construct, tagged with HA at the N-terminal, showed similar nuclear localisation (Supplementary Fig. 1g). These data raised the possibility that after cleavage, iR2-NT (but not iR2-CT) translocates to the nucleus.

To explore the characteristics of iRhom2 cleavage, we assessed the size of iR2-NT by expressing different deletion mutants of 2XHA tagged N-terminal domain. The N-terminal iRhom2 cleavage fragment was sized closest to iR2-1-403, which included predicted TMD1 and a few luminal amino acids (Fig. 1d), indicating that the cleavage occurs around the luminal border of TMD1, which would generate a fragment that remains membrane tethered. The nuclear localisation of the N-terminal domain was confirmed with immunofluorescent staining: iR2-1-374 and iR2-1-382, both of which lack full TMD1, were solely nuclear (Fig. 1e). In contrast, iR2-1-392 and iR2-1-403, both containing TMD1, showed both soluble nuclear and ER staining, including in the nuclear envelope; iR2-1-403 (Fig. 1f) and the naturally cleaved iR2-NT fragment (Supplementary Fig. 1h) both colocalised with the nuclear envelope marker SUN2.

Consistent with our experimental observations, we identified two conserved potential nuclear localisation signal (NLS) motifs in the iRhom2 N-terminus – one monopartite and one bipartite (Supplementary Fig. 1j). The traffic of membrane proteins with bulky cytoplasmic domains to the nuclear envelope is dependent on the presence of disordered regions and an NLS (Meinema et al., 2011; Mudumbi et al., 2020); in accordance with this, deletion of both NLS motifs from iR2-1-403 markedly reduced soluble nuclear staining (Fig. 1g).

Overall, these results lead us to conclude that proteolytic release of the iRhom2 N-terminal domain from the full-length ER-localised protein leads to its nuclear translocation. Note that the iR2-NT fragment contains TMD1 and thereby remains membrane tethered. Despite this, the diffuse nucleoplasmic staining of iR2-NT cleaved from full-length iRhom2 and the TMD-containing constructs iR2-1-392 and iR2-1-403 implies the existence of a secondary cleavage event that ultimately releases a soluble nuclear iRhom2 fragment into the nucleoplasm.

### iRhom2 is cleaved at the luminal juxtamembrane region of TMD1

The primary site of iRhom2 cleavage is predicted to be near the luminal end of TMD1 (Fig. 1d, 2a), the boundaries of which were further defined based on secondary structure and TMD prediction analyses (Supplementary Fig. 2a, b). Sequence alignment of iRhom1 and iRhom2 from mouse and human showed a highly conserved stretch of 8 amino acid residues (YGIAPVGF) in this region (Fig. 2a). Mutation to leucine of all (L8), or only the last 4 (L4), of these 8 amino acids abolished cleavage of iRhom2 (Fig. 2b), as did the mutation of PVGF to LVLF. In contrast, mutation of PV to AI only partially inhibited cleavage (Fig. 2b). We conclude that the PVGF motif is required for iRhom2 cleavage. Significantly, *Drosophila* iRhom does not contain the PVGF motif (PVGF being replaced by PIGI; Fig. 2c) and, consistent with the importance of the motif, we detected no cleavage of *Drosophila* iRhom (Fig. 2c). Furthermore, mutation of the PVGF in iRhom2 to the *Drosophila* sequence PIGI abrogated cleavage (Fig. 2d). The converse experiment, mutating *Drosophila* iRhom from PIGI to the mammalian sequence PVGF, caused *Drosophila* iRhom to be cleaved (Fig. 2d). Overall, these data confirm that the widely conserved (Supplementary Fig. 2c) PVGF motif in the luminal juxtamembrane region adjacent to TMD1 determines the cleavage of iRhom proteins.

**Fig. 2.**
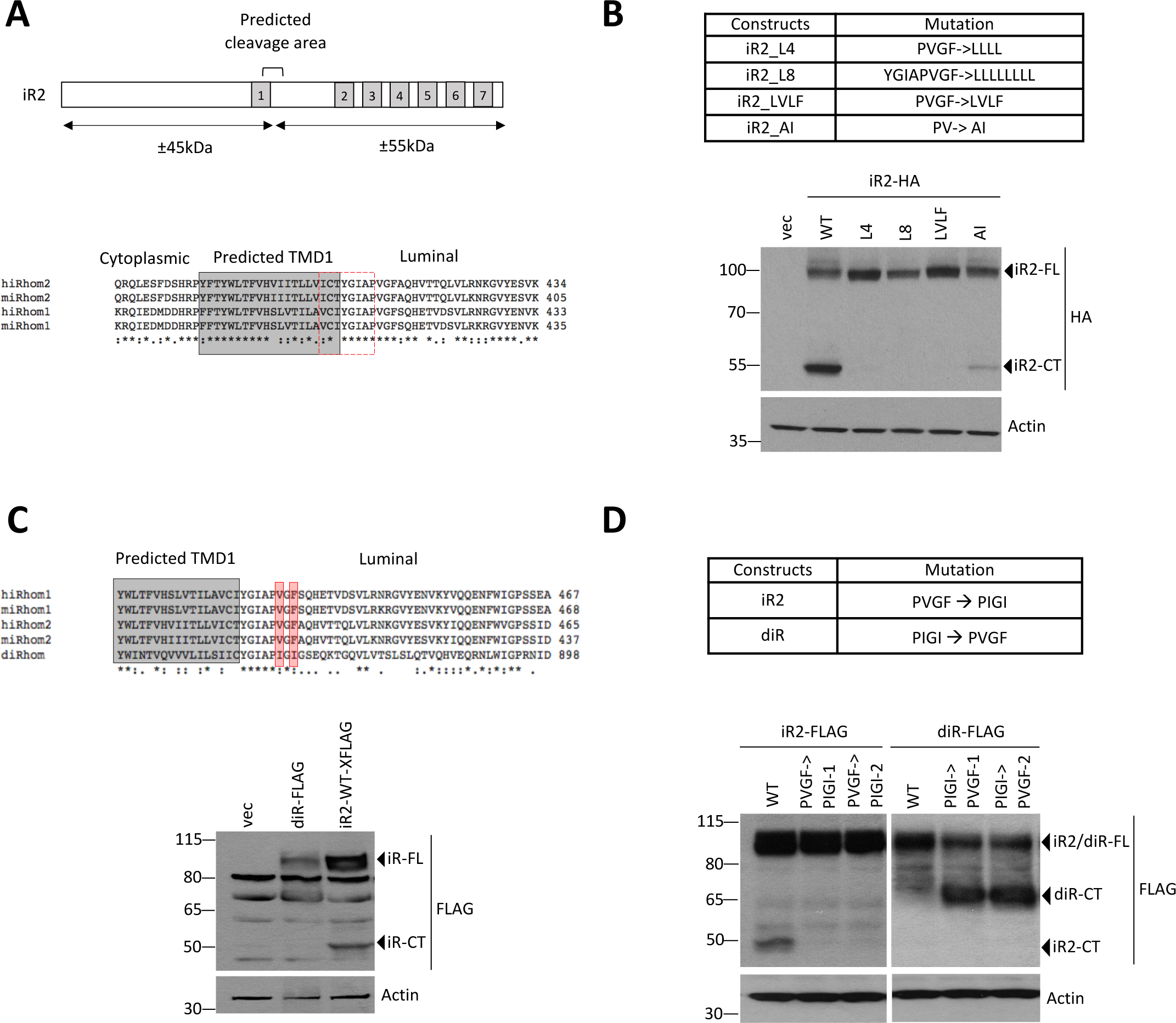
iRhom2 is cleaved at a conserved motif in the luminal domain. **a** Schematic showing predicted cleavage area of iRhom2 (top). Alignment of protein sequences of iRhom1 and iRhom2 from *Homo sapiens* (h) and *Mus musculus* (m). Boxed sequences are the predicted TMD1 (grey) and the highly conserved region juxtamembrane adjacent to TMD1 (dashed red). **b** Wild-type and iRhom2 mutans (indicated above) within the predicted cleavage region were analysed by immunoblotting in lysates from HEK293T cells transiently transfected for 36 h (bottom), using HA antibody. **c** Alignment of iRhom protein sequences from *Homo sapiens* (h) and *Mus musculus* (m) and *Drosophila melanogaster* (d). Boxed sequences are the predicted TMD1 (grey) and the two amino acids different in *Drosophila* are highlighted in red. iRhoms were analysed by immunoblotting in lysates from HEK293T cells transiently transfected for 36 h with *Drosophila* iRhom and human iRhom2 C-terminally tagged with 3XFLAG. **d** Levels of human iRhom2 and *Drosophila* iRhom mutated within PVGF and PIGI region (top) were analysed by immunoblotting in lysates from HEK293T cells transiently transfected for 36 h (bottom). 1 and 2 denote two different constructs for the same mutations.

We assessed whether uncleavable iRhom2 mutants can fulfil two well characterised functions: the destabilisation of EGF-like ligands (Zettl et al., 2011), and the maturation of ADAM17 (Adrain et al., 2012; McIlwain et al., 2012). All uncleavable mutants were able to downregulate EGF protein levels indistinguishably from wild-type iRhom2, implying that these iRhom2 mutants are functional (Supplementary Fig. 2d). ADAM17 maturation was also unaffected by the uncleavable mutant PVGF->PIGI, implying that this process, too, was not dependent on iRhom2 cleavage (Supplementary Fig. 2e). ADAM17 maturation was, however, inhibited by the other uncleavable iRhom2 mutants (AI, L4, L8) (Supplementary Fig. 2e). This apparently contradictory result is explained by co-immunoprecipitation assays, which show that those iRhom2 mutants defective in ADAM17 processing had markedly reduced binding between iRhom2 and ADAM17 (Supplementary Fig. 2f), whereas the PVGF->PIGI mutation binds normally. Overall, these results indicate that neither EGF degradation nor ADAM17 maturation depend on iRhom2 cleavage.

### Signal peptidase complex cleaves iRhom2

We sought to identify the protease responsible for primary iRhom2 cleavage. Structural prediction by AlphaFold agreed with TMD predictions that the PVGF motif is located within the ER lumen, immediately adjacent to TMD1 (Fig. 3a). Treatment with broad spectrum protease inhibitors against each of the four main categories of proteases (serine, cysteine, aspartic acid, and metallo-) did not significantly inhibit cleavage of iRhom2 (Supplementary Fig. 3a). One possibility we checked was RHBDL4, a rhomboid protease, located in the ER, and capable of cleaving within the luminal domains of its substrates (Fleig et al., 2012; Paschkowsky et al., 2016), but its knockdown had no effect on iRhom2 cleavage (Supplementary Fig. 3b).

**Fig. 3.**
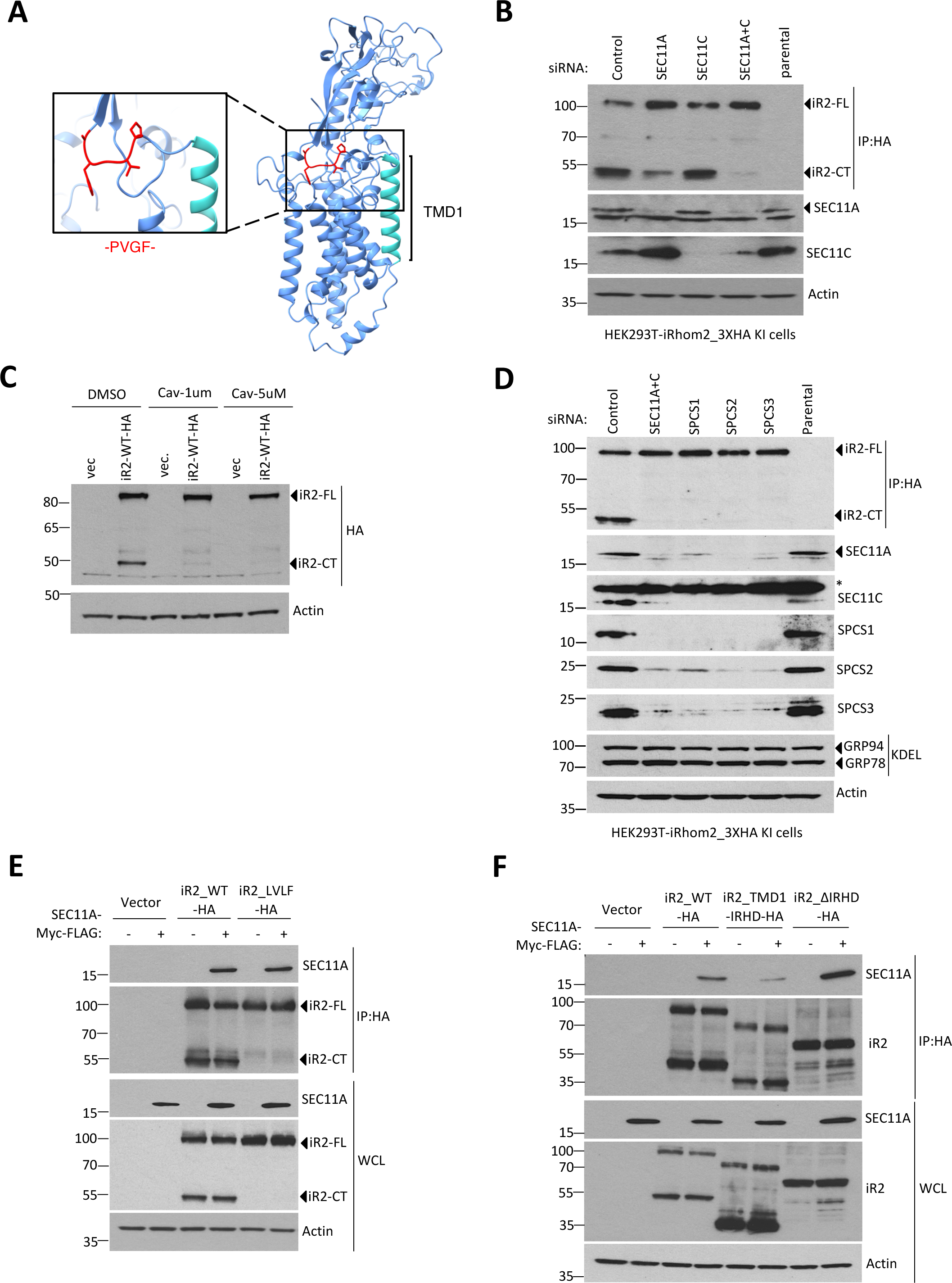
iRhom2 is cleaved by the signal peptidase complex. **a** Structural prediction for iRhom2 by AlphaFold. The region containing amino acids necessary for iRhom2 cleavage (PVGF) is highlighted in red. **b** Endogenously tagged iRhom2-HA was detected by immunoblotting in lysates from HEK293T cells transfected for 72 h with control siRNA or siRNA against the catalytic subunits SEC11A or SEC11B, or both combined. **c** C-terminally tagged iRhom2 under a TET-inducible promoter expressed for 18h was detected by immunoblotting in lysates from HEK293T cells in the presence of cavinafungin (Cav) at indicated concentrations. **d** Endogenously tagged iRhom2-HA was detected by immunoblotting in lysates from HEK293T cells transfected for 72 h with control siRNA or siRNA against the indicated components of the SPC. **e-f** iRhom2-HA immunoprecipitates SEC11A. Wild-type or mutant forms of iRhom2 (**e**-**f**) and SEC11A were analysed by immunoblotting in whole cell lysate (WCL) and immunoprecipitation (IP: HA) from HEK293T cells transiently transfected for 36 h with SEC11A-Myc-FLAG and iRhom2 constructs.

We have previously reported an iRhom2 interaction screen, in which one of the top hits was SEC11C (Kunzel et al., 2018), one of the two catalytic subunits of eukaryotic signal peptidase complex (SPC) (Liaci et al., 2021; Shelness and Blobel, 1990). Although SPC has a well-established function in removing canonical signal peptides from proteins entering the ER, it also catalyses the cleavage of multiple other signal peptide-like sequences (Hegde and Bernstein, 2006; Owji et al., 2018). Knockdown of SEC11C alone did not have any significant effect on cleavage of endogenous iRhom2 in HEK293T cells but silencing the alternative SPC catalytic subunit, SEC11A, had a slight effect (Fig. 3b). Most strikingly, the combined knockdown of SEC11A and SEC11C abolished cleavage of endogenous (Fig. 3b) or overexpressed iRhom2 (Supplementary Fig. 3c). Additionally, treatment of cells with cavinafungin, an inhibitor of SPC (Estoppey et al., 2017), blocked the cleavage of iRhom2 (Fig. 3c). In further support of the role of SPC, knockdown of any of the three essential accessory subunits (SPCS1, SPCS2, SPCS3) also blocked iRhom2 cleavage (Fig. 3d); note that knockdown of one subunit of SPC leads to depletion of the others, a phenomenon often seen in multi-subunit protein complexes (Pla-Prats and Thoma, 2022). In contrast, none of these treatments altered the expression of ER chaperones GRP78 or GRP94, indicating that SPC knockdown was not causing more general defects in ER homeostasis. Consistent with iRhom2 being a substrate of SPC, two signal peptide prediction algorithms (Hiller et al., 2004; Petersen et al., 2011) identified a potential SPC cleavage site in iRhom2, at the sequence PVGFA|QH, which matches the region we have experimentally determined to be required for cleavage (Supplementary Fig. 3d; Fig 2). As a negative control of this prediction, no cleavage sites were predicted around TMD1 of DERLIN1, another ER-localised rhomboid-like pseudoprotease (Supplementary Fig. 3d). Finally, the SEC11A catalytic subunit coimmunoprecipitated with both wild-type iRhom2 and the uncleavable LVLF mutant (Fig. 3e). Deletion of TMD2->7 of iRhom2 (iR2_TMD1_IRHD) but not the luminal IRHD (iR2_ΔIRHD) showed this interaction was dependent on the TMD domains of iRhom2 (Fig. 3f).

Since our data imply a secondary cleavage, to release the membrane tethered product of iRhom2 cleavage by SPC into the nucleoplasm (see above), we asked whether the intramembrane protease signal peptide peptidase (SPP) might be a responsible for the second cleavage. SPP cuts a number of released signal peptides after canonical SPC processing (Weihofen et al., 2002), although this appears not to be universal (Mentrup et al., 2017). Inhibition of SPP with (Z-LL)_2_ ketone had no effect on nuclear staining of iRhom2 (Supplementary Fig. 3e), indicating that SPP is not responsible for the formation of soluble iR2-NT.

### Nuclear iRhom2 modifies the cellular transcriptome

To ask whether the nuclear iRhom2 fragment has biological activity, we characterised the consequences of expressing the nuclear iR2-1-374 fragment (Fig. 1e). We generated HEK293T cells stably expressing HA-tagged iR2-1-374, and biochemical fractionation experiments detected iR2-1-374 in both soluble (S4) and insoluble, chromatin containing (P4) nuclear fractions (Supplementary Fig. 4a). The soluble (S4) fraction was increased by MNASE-mediated DNA digestion (Supplementary Fig. 4a), similar to control Histone H3 protein, indicating that nuclear iRhom2 associates with chromatin. We also generated HEK293T cells stably expressing HA-tagged iR2-1-374 under a tetracycline inducible promoter (Supplementary Fig. 4b), and observed that iR2-1-374 co-immunoprecipitated with the B1 subunit of RNA Polymerase II (POL2), and transcription factor IID (TFIID) (Supplementary Fig. 4c), both integral components of the eukaryotic gene transcription complex (Roeder, 2019). To test the implication that nuclear iR2-1-374 might therefore influence gene expression, using RNA-seq, we examined the cellular transcriptome at 3h and 6h after induction. The expression of 1233 and 1280 genes were significantly changed (adjusted p-value <0.05) at 3h and 6h respectively (Fig. 4a). At 3h induction, 404 genes were up-regulated, and 829 genes were down-regulated. At 6h induction, 382 genes were up-regulated, and 898 genes were down-regulated (Fig. 4a). A complete list of differentially expressed genes is provided in Supplementary Table 1.

**Fig 4.**
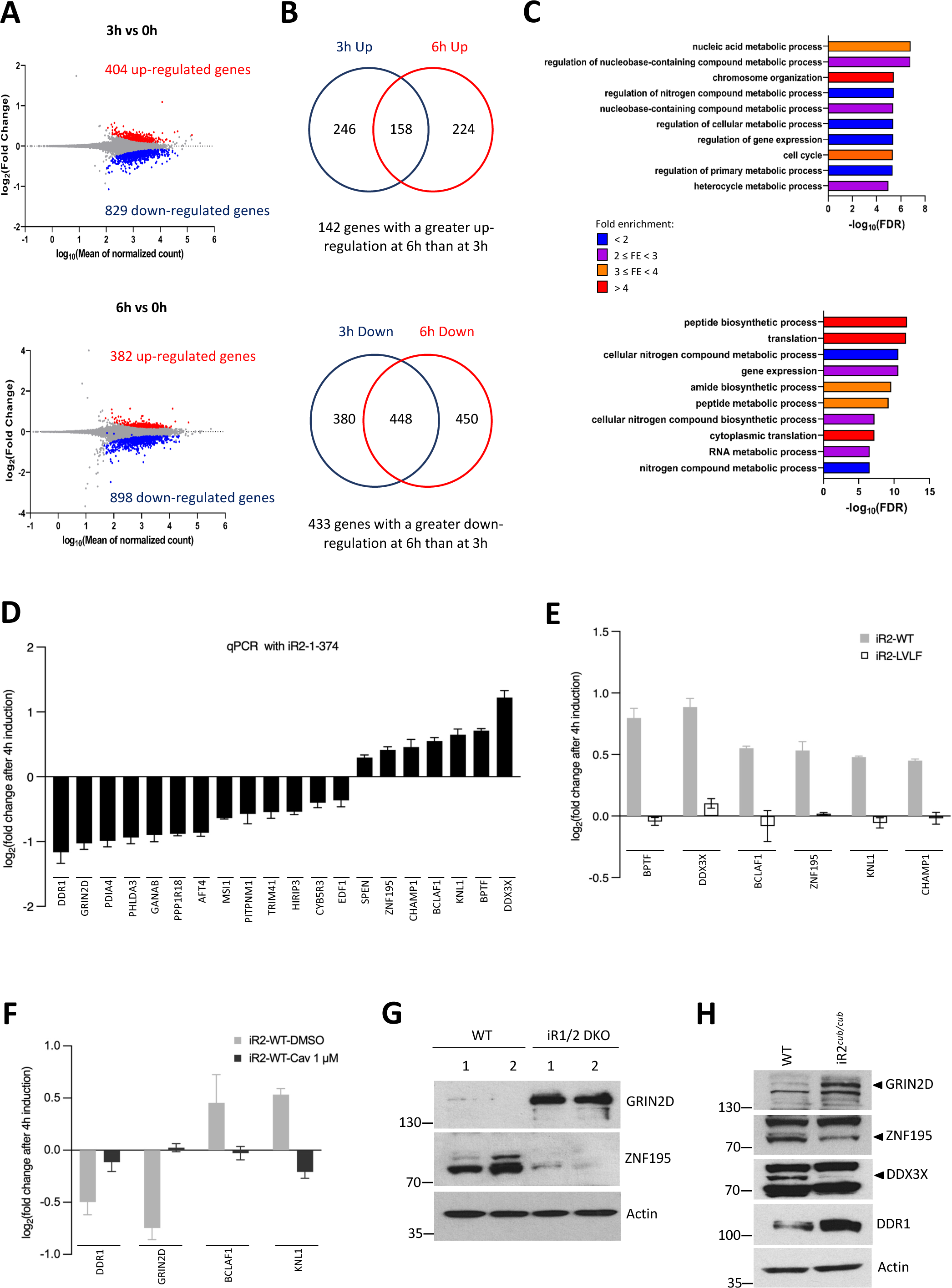
Nuclear iRhom2 induces gene expression changes. **a** MA plot showing differentially expressed genes (upregulated in red, downregulated in blue) in HEK293T cells expressing iR2-1-374 under a TET-inducible promoter expressed for 3 h or 6 h against the 0 h control. n=3, adjusted p-value < 0.05. **b** Venn diagrams showing overlap between significantly upregulated or downregulated genes at 3 h and 6 h of iR2-1-374 induction. Genes with a greater downregulation (n=433) or a greater upregulation (n=142) at 6 h than at 3 h of expression were considered as more likely targets of nuclear iRhom2. **c** Bar graphs showing summary of the Gene Ontology (GO) enrichment analysis on the specific set of upregulated (n=142) or downregulated (n=433) genes, categorised as likely targets of nuclear iRhom2. FDR: False discovery rate. Different colours denote range of fold enrichment (FE). **d** Graph showing validation of 20 potential target genes from RNA-seq data by quantitative RT-PCR (qPCR) in HEK293T cells expressing inducible iR2-1-374 for 4 h. Data presented as Log_2_ fold change relative to WT cells as mean ± SEM. **e-f** Graphs showing validation of selected target genes by qPCR in HEK293T cells expressing for 4 h either inducible wild-type (iR2-WT) or uncleavable iRhom2 (iR2-LVLF) (**e**) or inducible wild-type iRhom2 in the presence of Cavinafungin (1 µM) (**f**). Data presented as Log_2_ fold change relative to WT cells as mean ± SEM. **g-h** Protein levels of selected target genes of nuclear iRhom2 were analysed by immunoblotting in cell lysates from two independent pairs of wild-type and iRhom1/2 DKO (g) or iRhom2^cub/cub^ (h) mouse embryonic fibroblasts (MEFs) with indicated antibodies. qPCR and immunoblots data are representative of 2-3 independent experiments.

We analysed the genes that were either up-regulated (158 genes), or down-regulated (448 genes) at both 3h and 6h time points, and which showed a greater difference at 6h than at 3h (Fig. 4b). Using Gene Set Enrichment Analysis (GSEA) (Subramanian et al., 2005), we identified nucleic acid metabolism, chromosome organisation, regulation of gene expression and cell cycle as the top processes associated with genes up-regulated by iR2-1-374 (Fig. 4c, Supplementary Table 2), and peptide biosynthesis, translation and RNA metabolism as the most prominent down-regulated processes (Fig. 4c, Supplementary Table 2).

RT-qPCR validation of 20 differentially regulated genes confirmed that 7 were significantly upregulated by iR2-1-374, including *KNL1*, *BCLAF1*, *BPTF*, *DDX3X* and *ZNF195*, and 13 were significantly down-regulated, including *GRIN2D*, *DDR1*, *ATF4*, *GANAB*, *PHLDA3* and *PDIA4* (Fig. 4d). Notably, inducing expression of the uncleavable (LVLF) mutant of iRhom2 in HEK293 cells (Supplementary Fig. 4d), led to no alteration of expression of validated target genes (Fig. 4e). Similarly, expression of wild-type iRhom2 in the presence of cavinafungin to inhibit SPC-mediated cleavage (Supplementary Fig. 4e) prevented alteration of target gene expression (Fig. 4f). Both these results confirm the requirement for iRhom2 cleavage for changes in gene expression of target genes.

We also asked whether the transcriptional regulation of these genes by iR2-1-374 led to changes at the protein level. We selected four genes which showed most consistent qPCR changes caused by wild-type iRhom2 and iR2-1-374 and did this experiment two ways: (1) comparing two independently-derived lines of wild-type and iRhom1/2 double knockout (iR1/2DKO) mouse embryonic fibroblasts (MEFs) (Christova et al., 2013); and (2) comparing wild-type and iRhom2*^cub/cub^* MEFs, the latter containing a mutation in endogenous mouse iRhom2 that removes most of the cytoplasmic N-terminal domain(Hosur et al., 2014). Levels of GRIN2D and ZNF195 proteins were increased and decreased respectively in absence of iRhom1 and iRhom2 (Fig. 4g). Conversely, iRhom2*^cub/cub^* cells, which lack endogenous nuclear iRhom2, showed upregulation of GRIN2D and DDR1, and downregulation of ZNF195 and DDX3X (Fig. 4h). Together, these results are consistent with the transcriptional changes observed by RNA-Seq and RT-qPCR; they also indicate that the regulation of some target genes are conserved between human and mouse cells.

### Nuclear iRhom2 expression is enhanced in human skin pathologies

Beyond a well characterised role in inflammatory signalling in macrophages, one of the main sites of iRhom2 expression is skin (Christova et al., 2013). Moreover, mutations in the iRhom2 N-terminal domain cause skin pathologies in the inherited disease tylosis with oesophageal cancer (TOC) (Blaydon et al., 2012; Brooke et al., 2014). It was therefore striking to observe that immunohistochemical staining of patient normal skin samples showed detectable expression of iRhom2 in the nuclei, which was elevated in the suprabasal and basal layers of lesional (TOC) interfollicular skin, as well as in the epidermis of psoriatic skin (Fig. 5a). Nuclear iRhom2 was also observed in immune cell populations residing within the dermis, particularly in lesional psoriatic skin biopsies (Fig. 5a).

**Fig 5.**
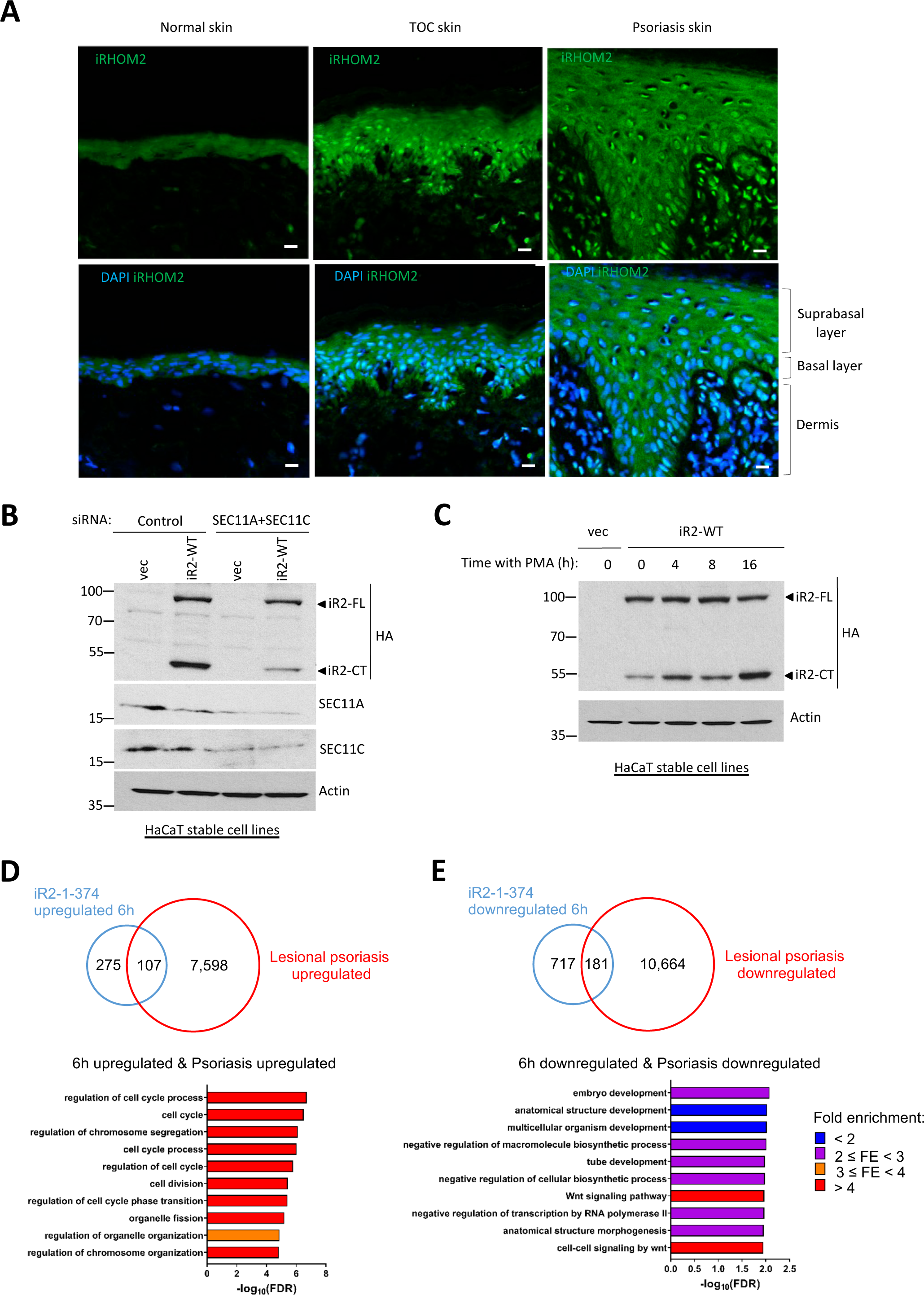
Human skin diseases show high levels of nuclear iRhom2. **a** Levels of iRhom2 expression were determined by immunohistochemistry in the basal, suprabasal and dermis layers in both lesional TOC interfollicular and psoriatic skin epidermis compared to control interfollicular skin. Tissues were stained for iRhom2 (green) and DAPI (blue). Scale bar: 20 *μ*m. **b-c** iRhom2 was detected by immunoblotting in lysates from HaCaT cells stably expressing iRhom2-3XHA after transfection with either control siRNA or siRNA against SEC11A and SEC11C for 72 h before harvest (**b**), or after treatment with 200 nM PMA for indicated time intervals (**c**). **d-e** Venn diagrams showing overlap between differentially expressed genes found in a published RNA-Seq dataset of lesional psoriasis and upregulated (**d**) or downregulated (**e**) at 6 h induction of iR2-1-374 (top). Bar graphs showing summary of the GO enrichment analysis on the set of common upregulated (n=107) or downregulated (n=181) genes between lesional psoriasis and iR2-1-374. FDR: False discovery rate. Different colours denote range of fold enrichment (FE).

To explore the proposed nuclear signalling of iRhom2 in skin, we stably expressed iRhom2 in HaCaT cells, a human epidermal keratinocyte line. As in HEK293T cells (Fig. 3), knockdown of the catalytic subunits of signal peptidase complex (SEC11A and SEC11C) strongly inhibited the cleavage of iRhom2 (Fig. 5b). Psoriasis is characterised by abnormal proliferation and differentiation of keratinocytes, combined with chronic inflammation (Lowes et al., 2014) and treatment with the phorbol ester PMA is commonly used to model the epidermal thickening and dermal inflammation of psoriatic lesions (Chang et al., 2020; Zhang et al., 2013). Significantly, PMA treatment of iRhom2-expressing HaCaT and HEK293 cells led to an increase in the cleavage of iRhom2 (Fig. 5c; supplementary Fig. 5a). This enhanced cleavage was significantly abrogated by knockdown of SPC catalytic subunits (Supplementary Fig. 5b), indicating that PMA-induced cleavage of iRhom2 is mediated by SPC.

Analysis of a published dataset of high-depth RNA-Seq of the skin transcriptome from healthy controls, non-lesional psoriatic, and lesional psoriatic patients (Tsoi et al., 2019) revealed increased expression of iRhom2 (but not iRhom1), as well as SEC11C, SPCS1 and SPCS2 in lesional psoriasis tissue compared to healthy skin (Supplementary Fig. 5c). Similar upregulation was observed between non-lesional and lesional psoriasis (Supplementary Fig. 5d). We also found significant overlap in gene expression changes between lesional psoriasis and cells expressing iR2-1-374: 107 upregulated genes (Fig. 5d) and 181 downregulated genes were common to both lesional psoriasis and cells expressing iR2-1-374 for 6h (Fig. 5e); at 3h, 104 genes in common were upregulated (Supplementary Fig. 5e), and 149 genes in common downregulated (Supplementary Fig. 5f). A complete list of differentially expressed genes is provided in Supplementary Table 3. Gene ontology analysis revealed in both sets a significant enrichment of genes regulating processes including cell cycle, chromosome segregation, Wnt signalling pathway (at 6h; Fig. 5d, e), and nucleic acid metabolism, cytoplasmic translation and cell development (at 3h; Supplementary Fig. 5e,f). A ranked list of all biological processes is provided in Supplementary Table 4. Together these results indicate the biological significance of nuclear iRhom2 expression in the skin, and suggest that elevated levels may contribute to the pathology of skin diseases.

## DISCUSSION

iRhoms, which are primarily located in the ER and the plasma membrane, are the best studied non-protease members of the rhomboid-like superfamily (Dulloo et al., 2019; Lemberg and Adrain, 2016). Although the biological role of rhomboid-like proteins is varied, a common functional theme has been their specific recognition of TMDs in substrates and clients. In this work, we have uncovered an iRhom-mediated signalling pathway, unrelated to this core function of membrane protein regulation. It is mediated by the proteolytic release of the iRhom N-terminal domain, which then translocates to the nucleus, where it regulates gene expression. By this mechanism, iRhoms now join a select group of membrane proteins with a secondary nuclear function triggered by proteolytic release of intracellular domains. The best-known examples include Notch, SREBP (sterol regulatory element-binding protein), and the ATF6 branch of the unfolded protein response (De Strooper et al., 1999; Sakai et al., 1996; Ye et al., 2000). Cytoplasmic domains of each of these proteins are released from cellular membranes, to translocate to the nucleus where they trigger a transcriptional response.

In each of those previously discovered examples, targets are cleaved by an intramembrane protease, which cuts its substrate TMD in a position that generates a fragment with too short a hydrophobic helix to be retained in the membrane. In the case of SPC cleavage of iRhom2, which we believe to be the first report of a mammalian membrane tethered protein being converted to a nuclear signal by SPC, there must be a secondary cleavage to release the cleaved N-terminal domain from its membrane anchoring TMD. Indeed, we see that iRhom2 in the nucleus exists in two distinct sub-locations: membrane tethered in the nuclear envelope, and diffusely located in the nucleoplasm. As expected, when the N-terminal domain with no TMD is expressed, it is solely located in the nucleoplasm. We do not know the identity of the protease responsible for the secondary cleavage, reflecting wider uncertainty about the fate of canonical signal peptides cleaved by SPC. Some are degraded by the intramembrane aspartyl protease, signal peptide peptidase (SPP), which was originally named to reflect the view that it was the protease that degrades all signal peptides (Weihofen et al., 2002). More recently, however, it has become much less clear that SPP has a general role, and the fate of most signal peptides after SPC cleavage remains uncertain (Mentrup et al., 2017). In the case of iRhom2, we have experimentally ruled out SPP as the secondary protease that releases the soluble form of the N-terminus.

Signal peptidase complex is primarily responsible for the removal of signal peptides from proteins entering the ER (Jackson and Blobel, 1977; Liaci et al., 2021). Signal peptides are typically 15-30 amino acids long, and reside in the first 30 amino acids of the coding sequence. Despite being in a TMD nearly 400 amino acids from the N-terminus of the protein, the SPC cleavage site of iRhom2 broadly resembles the normal determinants of signal peptide cleavage (von Heijne, 1983, 1984): two positively charged conserved amino acids (H and R) (the n-region), immediately preceding TMD1 (the h-region), followed by uncharged conserved amino acid segment (YGIAPVGF) (the c-region). At the predicted cleavage site of iRhom2 (PVGFA|Q), residues at positions -1 and -3 are Ala and Gly respectively and, together with a Gln residue at the +1 position, these align with the consensus amino acids observed in eukaryotic signal peptidase cleavage sites (Choo and Ranganathan, 2008). Accordingly, the iRhom2 cleavage site is recognised by signal peptide identification algorithms (as long as they do not incorporate a penalty for distance from the N-terminus). Interestingly, the SPC substrate-determining amino acids are well conserved in iRhoms across many species, although a few (chicken, some fish) have variations that might impair cleavage. In contrast, they are absent in the single *Drosophila* iRhom. Consistent with this prediction, we have shown *Drosophila* iRhom to be uncleaved (and uncleavable) by SPC, unless the relevant amino acids are mutated to the human sequence. Other non-canonical cleavage of TMDs by SPC has been previously reported. For example, SPC removes 83 residues from the *Drosophila* cell surface protein Crumbs (Kilic et al., 2010), 37 residues from the human cytomegalovirus protein UL40 (Ulbrecht et al., 2000), and 135 residues from the canine distemper virus fusion glycoprotein F0 (von Messling and Cattaneo, 2002). To our knowledge, the N-terminal domain of iRhom2 is the longest N-terminal fragment from a mammalian protein to be reported as cleaved by SPC.

It is essential that iRhoms are only partially cleaved by SPC, because other iRhom functions, like ADAM17 activation and response to ER stress, require full-length iRhom2. The mechanistic basis for this partial SPC cleavage of iRhom2, and how it is regulated, remains unknown but it is notable that the timing and efficiency of signal sequence cleavage does vary for different SPC substrates. In the case of the HIV-1 gp160 envelope protein, for example, slow and inefficient signal sequence cleavage provides a checkpoint mechanism to ensure full folding and maturation has occurred prior to onward trafficking from the ER (Snapp et al., 2017). The flaviviridae family also take advantage of slow processing by endogenous SPC, to ensure a correct sequence of cleavages of the viral polyprotein – necessary for efficient virus particle assembly and propagation (Alzahrani et al., 2020). In both these cases, the cause of inefficient or slow processing has been mapped to non-canonical sequences in the SPC recognition sequence, so it will be interesting in the future to explore more widely the regulation and precise determinants of iRhom cleavage by SPC.

The regulation of the cellular transcriptome by nuclear iRhom2 is likely to be indirect as it does not have any discernible features of a typical transcription factor (e.g., transactivation or DNA-binding domains). Indeed, our observations that nuclear iRhom2 can bind to chromatin, and interact with RNA Polymerase II and TFIID, components of the eukaryotic transcription complex (Roeder, 2019), indicate that its regulation of gene expression is likely as part of a transcription activator or repressor complex. It would seem most likely that the soluble form of nuclear iRhom2-NT is responsible for its transcriptional regulatory function, but we note that it is also possible that the membrane tethered nuclear iRhom2-NT could play a more direct role in effecting gene expression changes. Nuclear envelope tethered proteins mostly act through interaction with chromatin associated proteins (Czapiewski et al., 2016), LAP2β and MAN1 can bind directly to DNA (Cai et al., 2001; Caputo et al., 2006). MAN1 can also act as a transcription scavenger (Lin et al., 2005; Pan et al., 2005), sequestering R-Smads, crucial regulators of transforming growth factor-β (TGFβ), bone morphogenic protein (BMP) and activin signalling (Massague et al., 2005). Note, however, that despite this molecular question of the relative significance of membrane tethered and nucleoplasmic iRhom2-NT, the transcriptional effects of iRhom2 are strictly SPC-dependent.

Dominant iRhom2 mutations are the cause of the inherited syndrome tylosis with oesophageal cancer (TOC), which is characterised by palmoplantar keratoderma, oral precursor lesions, and a high lifetime risk of oesophageal cancer (Blaydon et al., 2012). iRhom2^TOC^ mutants show upregulated shedding of EGF ligands by ADAM17 (Brooke et al., 2014; Maney et al., 2015), which is proposed to contribute to disease pathology. Our discovery of a parallel nuclear function of the iRhom2 N-terminal domain, indicates that iRhom2-dependent changes to gene expression may also contribute to pathogenesis. Consistent with this idea, we show nuclear localisation of iRhom2 in normal human skin, which is significantly elevated in TOC and lesional psoriatic skin, indicating a possible association between levels of nuclear iRhom2 and disease. Although this potential pathogenic mechanism needs further exploration, it is striking that both TOC and psoriasis exhibit epidermal keratinocyte hyperplasia, and that epidermal thickness has been associated with iRhom2 expression in both mice and humans (Blaydon et al., 2012; Hosur et al., 2017a; Hosur et al., 2017b; Maruthappu et al., 2017). It is also notable that both iRhom2 and SPC subunits are upregulated in psoriasis in our analysis of a published RNA-Seq dataset (Tsoi et al., 2019). The relevance of iRhom2 to the pathogenesis of psoriasis is also consistent with the overlap of genes differentially regulated by nuclear iRhom2, and those in lesional psoriasis. Notably, GO analysis indicates that one of the main biological processes predicted to be affected in both data sets is the cell cycle. Psoriatic skin lesions develop following increased mitosis of keratinocytes, leading to incomplete cornification and a poorly adherent stratum corneum (Griffiths and Barker, 2007). Several studies also noted that the mitotic cell cycle was among the most significant process differentially regulated between normal and lesional psoriasis skin tissues (Pasquali et al., 2019; Tsoi et al., 2019; Xie et al., 2014). Nevertheless, we want to emphasise that there is much to learn about the role of iRhom2 in skin pathologies. For example, as well as increased cell division, psoriasis is also characterised by acute infiltration of inflammatory immune cells including dendritic cells, macrophages and T cells, which induce the release of chemokines and cytokines, including TNF (Nestle et al., 2009). iRhom2 is highly expressed in immune cells and, through its central role in ADAM17 activation, controls TNF secretion (Adrain et al., 2012; Al-Salihi and Lang, 2020; McIlwain et al., 2012). Overall, therefore, it seems likely that the pathogenesis of skin disease associated with iRhom2 may combine aspects of both inflammatory signalling and nuclear functions.

Finally, iRhoms represent an interesting example of the still rather poorly explored phenomenon of pseudoenzymes and their potential biological significance (Adrain and Freeman, 2012; Ribeiro et al., 2019). Specifically, they are pseudoproteases, having lost through evolution the proteolytic activity of their more ancient cousins, the rhomboid intramembrane serine proteases. Until now, all known iRhom function has been apparently related to the proposed core function of members of the rhomboid-like superfamily, namely the specific recognition of TMDs and regulation of transmembrane proteins. Not only does this work expand our specific understanding of iRhom function, and illustrate the modular nature of rhomboid-like proteins, it also highlights how evolution can build completely new functions into pseudoenzymes, increasing functional divergence from their ancestral enzymes.

## EXPERIMENTAL PROCEDURES

### Molecular cloning and plasmids

Wild-type iRhom1, iRhom2 and mutants iRhom2_ΔIRHD and iRhom2_TMD1-IRHD plasmids in pEGFP-N1 vector with a C-terminal 3XHA tag have been previously described (Adrain et al., 2012; Dulloo et al., 2022). Wild-type iRhom2 and truncated mutants of iRhom2 (1-374, 1-382, 1-392,1-403,1-403_ΔmbNLS were amplified from iRhom2 cDNA by PCR and cloned with a N-terminal 2XHA tag into pEGFP-N1 mammalian expression vector (without EGFP protein). Wild-type iRhom2 and *Drosophila* iRhom were cloned by PCR into pEGFP-N1 vector with a C-terminal 3XFLAG tag. GFP-iRhom2-HA construct was generated by cloning of iRhom2-3XHA into pEGFP-N1 mammalian expression vector, in frame with EGFP protein at the N-terminal. Wild-type iRhom2 and iRhom2-1-374 were also cloned by PCR into pM6P.Blasticidin (for retroviral infection) and into pLVX-TetOne-Puro vector (Clontech, #631849) (for lentiviral infection). SEC11A-Myc-FLAG was obtained from OriGene Technologies (#RC204971).

All cloning PCR were done using Q5 High-Fidelity DNA Polymerase (New England Biolabs, #M0491S) and InFusion HD cloning according to manufacturer’s protocol (Takara Bio, #639649). Site-directed mutagenesis (SDM) was done by PCR method and DpnI digestion. All constructs were verified by Sanger sequencing (Source Bioscience, Oxford, UK).

### Human samples

Skin biopsies were obtained with written informed consent and approved by the Barts Health NHS Trust ethics committee (IRAS Project ID: 08/H1102/73). The study protocol conforms to the ethical guidelines of the Declaration of Helsinki.

### Mouse studies

Organs from wild-type and iRhom2 knockout mice previously generated (Adrain et al., 2012; Christova et al., 2013), were collected from sacrificed animals and stored on dry ice or at −80°C. Tissues were lysed in Triton X-100 RIPA buffer (1% Triton X-100, 150 mM NaCl, 50 mM Tris-HCl (pH 7.5), 0.1% SDS, 0.5% sodium deoxycholate) supplemented with EDTA-free protease inhibitor mix using a tissue homogeniser (Omni International). Lysates were cleared from cell debris by centrifugation (20,000 g, 4°C, 10 min) and used for immunoblotting. All procedures on mice were conducted in accordance with the UK Scientific Procedures Act (1986) under a project license (PPL) authorized by the UK Home Office Animal Procedures Committee, project licenses 80/2584 and 30/2306, and approved by the Sir William Dunn School of Pathology Local Ethical Review Committee.

### Cell culture and treatment

Mouse embryonic fibroblasts (MEFs) were isolated from Rhbdf1^−/−^/Rhbdf2^−/−^ (referred to as iRhom1/2 DKO) E13.5 embryos and wild-type C57BL/6J (RRID: IMSR_JAX:000664) controls and immortalised by lentiviral transduction with SV40 large T antigen as previously described(Adrain et al., 2012; Christova et al., 2013). MEFs cells from iRhom2^cub/cub^ and wild-type littermates have previously been described (Siggs et al., 2014). Human embryonic kidney (HEK293T) (#CRL-3216) and human epidermal keratinocyte (HaCaT) (#PCS-200-011) cells were obtained from ATCC, and HEK293T-iRhom1/2 DKO cells stably expressing inducible HA-iRhom2 has previously been described (Dulloo et al., 2022). HEK293T, HaCaT and MEFs cells were all cultured in high-glucose DMEM (Sigma-Aldrich, #D6429) supplemented with 10% fetal bovine serum (FBS) (Thermo Fisher Scientific, #10500064) and 5 mM glutamine (Gibco, #11539876) at 37°C with 5% CO_2_.

The following drugs were used: Cycloheximide (Sigma Aldrich, #C4859), Doxycycline (Sigma Aldrich, #D9891), MG-132 (Sigma Aldrich, #474791), Chloroquine (Sigma Aldrich, #C6628), 3-MA (Sigma Aldrich, #189490), E-64d (Sigma Aldrich, #E8640), Pepstatin A (Sigma Aldrich, #P5318), AEBSF (Sigma Aldrich, #101500), 3,4-Dichloroisocoumarin (3,4 DCI) (Sigma Aldrich, #D7910), phorbol 12-myristate 13-acetate (PMA) (Sigma Aldrich, #P8139), Cavinafungin (gift from Martin Spiess), Z-LL_2_ Ketone (Merck Life Science, #SML1442). All drug concentrations are indicated in figure legends and in respective methods sections. For cycloheximide chase assay, HEK293T cells transfected with indicated plasmids after 24h were treated with 100 µg/ml and harvested at indicated time points.

### Biochemical fractionation and micrococcal nuclease (MNase) treatment

Process was performed as previously described (Wysocka et al., 2001). 2 × 10^7^ cells were harvested and resuspended in 500 μl buffer A (10 mM HEPES [pH 7.9], 10 mM KCl, 1.5 mM MgCl_2_, 0.34 M sucrose, 10% glycerol, 1 mM dithiothreitol, and protease inhibitor cocktail). Triton X-100 was added (0.1% final concentration), the cells were incubated on ice for 8 min, and nuclei (fraction P1) were collected by centrifugation (5 min, 1,300 × g, 4°C). The supernatant (fraction S1) was clarified by high-speed centrifugation (5 min, 20,000 × g, 4°C), and the supernatant (fraction S2) was collected. The P1 nuclei were washed once in buffer A and lysed for 30 min in buffer B (3 mM EDTA, 0.2 mM EGTA, 1 mM dithiothreitol, and protease inhibitor cocktail), and insoluble chromatin (fraction P3) and soluble (fraction S3) fractions were separated by centrifugation (5 min, 1,700 × g, 4°C). The P3 fraction was washed once with buffer B and was resuspended in a solution containing 10 mM Tris, 10 mM KCl, and 1 mM CaCl_2_, with or without 1 U of MNase (Sigma Aldrich, #3755). After 15 min incubation at 37°C, the reaction was stopped with EGTA (1 mM final concentration).

Soluble and insoluble components were then separated by centrifugation (5 min, 1,700 × g, 4°C). The pellet was resuspended in in 2× SDS sample buffer and sonicated before incubation at 65°C for 15 min and used for immunoblotting.

### Transfection and transduction of cell lines

HEK293T cells transiently transfected with DNA in OptiMEM (Gibco, #10149832) using FuGENE HD (Promega, #E2312) and protein expression was analysed 24–48 h post transfection.

For knockdown experiments, siRNA was transfected using Lipofectamine RNAiMax (Thermo Fischer Scientific, #13778075) according to the manufacturer’s protocol. ON-TARGETplus SMARTpool human siRNA (Dharmacon) for SEC11A (#L-006038-00-0005), SEC11C (#L-046035-01-0005), SPCS1(#L-020577-00-0005), SPCS2(#L-020897-00-0005), SPCS3 (#L-010124-00-0005), and RHBDF2 (#HSS128594, #HSS128595, ThermoFisher Scientific), RHBDL4 (#HSS125697, #HSS125698, ThermoFisher Scientific) and non-targeting siRNA control (Dharmacon: #D-001206-13-50, ThermoFisher Scientific: Stealth RNAi™ siRNA Negative Control: #12935300) were used. Protein expression was analysed 72 h post siRNA transfection.

HEK293T and HaCaT cells stably expressing wild-type iRhom2-3XHA or iRhom2-1-374-3XHA were generated by retroviral transduction using pM6P retroviral constructs as previously described(Dulloo et al., 2022; Kunzel et al., 2018). In brief, HEK293T cells were transfected with indicated gene expressed in pM6P constructs together with packaging plasmid pCL.10A1. Viral supernatants for individual constructs were harvested after 48 h, cleared by centrifugation at 20,000 x g for 20mins, and co-incubated with HEK293T or HaCaT cells in the presence of 5 μg/ml polybrene and cells were selected with 10 μg/ml blasticidin (Sigma Aldrich, #15205). HEK293T expressing either iRhom2-1-374 or wild-type iRhom2 under TET-inducible cells were generated by lentiviral infection using gene cloned into pLVX-TetOne vector (Clontech, #631849). Methodology is similar to retroviral transduction, with exception of packaging vectors (pCMV-VSV-G and pCMV-dR8.91) and selected with 2 μg/ml puromycin (Thermo Fischer Scientific, #A1113803).

### Antibodies

For immunoblotting and co-immunoprecipitation: Actin (Santa Cruz, #sc-47778; 1:5000), FLAG-HRP (Sigma Aldrich, #A8592; 1:4000), HA-HRP (Roche, #11867423001; 1:2000), KDEL (AbCam, #ab12223; 1:2000), iRhom2-NT-specific ((Adrain et al., 2012); 1:500), Myc (Abcam, #ab9132; 1:2000), SEC11A (Proteintech, #14753-1-AP; 1:500), SEC11C (Novus Biologicals, NBP1-80774; 1:500), SPCS1 (Proteintech, #11847-1-AP; 1:500), SPCS2 (Merck Life Science, #HPA013386; 1:500), SPCS3 (Santa Cruz, sc-377334; 1:500), RHBDL4 ((Fleig et al., 2012); 1:1000), POL2 (MBL International, MABI0601; 1:1000), TFIID (Santa Cruz, #sc-273; 1:1000), Histone H3 (Cell Signaling, #4499T; 1:4000), GRIN2D (Novus Biologicals, NBP2-94573; 1:1000), ZNF195 (Novus Biologicals, NBP2-93054; 1:1000), DDR1 (Cell Signaling, #5583T ; 1:1000), DDX3X (Cell Signaling, #3189S; 1:1000), ADAM17 (AbCam, #ab39162; 1:2000).

For immunofluorescence: DAPI (Thermo Fischer, #D1306; 1 µg/ml), HA (Cell Signaling Technology, #3724; 1:500), HA.11 (Enzo Life Sciences, #ABS120-0200; 1:500), BAP31 (Enzo Life Sciences, #ALX-804-601-C100; 1:250), GFP (AbCam, #ab13970; 1:500), RHBDF2 (Sigma-Aldrich, #SAB1304414), SUN2 (Atlas Antibodies, HPA001209; 1:250)

### SDS-PAGE and immunoblotting

Cell lysates were denatured at 65°C for 15 min and ran either on 4-12 % NuPAGE™ Bis-Tris gels or 8-16 % Tris-Glycine Novex™ WedgeWell™ (Thermo Fischer Scientific, #NP0321, #XP08160) in MOPS or Tris-Glycine running buffer respectively. PageRuler™ Plus Prestained Protein Ladder (Thermo Fischer Scientific, #26620) was used for protein molecular weight marker. Note this ladder runs differently on Bis-Tris and Tri-Glycine gels, resulting in different molecular weights according to manufacturer. Both types of gel were used throughout study and gels were transferred onto polyvinylidene difluoride (PVDF) membranes (Millipore, #IPVH85R). The membrane was blocked in 5 % milk-TBST (150 mM NaCl, 10 mM Tris-HCl pH 7.5, 0.05 % Tween 20, 5 % dry milk powder) before incubation with indicated primary and species-specific HRP-coupled secondary antibodies. All primary antibodies were made in 5% BSA-TBST except for HRP-conjugated antibodies. Band visualisation was achieved with Amersham Enhanced Chemiluminescence (GE Healthcare, #RPN2106)) or SuperSignal™ West Pico PLUS Chemiluminescent Substrate (Thermo Fisher Scientific, #34577) using X-ray film. Quantification of blots was done using Fiji (Image J).

### Co-immunoprecipitation

HEK293T cells were transfected in 6-cm plates with indicated plasmids for 36-48 h before harvest. MEFs cells stably expressing iRhom2 were also processed according to the following steps. Cells were washed with ice-cold PBS and then lysed on ice in Triton X-100 lysis buffer (1 % Triton X-100, 200 mM NaCl, 50 mM Tris-HCl pH 7.4) supplemented with cOmplete™, EDTA-free Protease Inhibitor Cocktail (Merck, #4693132001). Cell lysates were cleared by centrifugation at 21,000 x g for 20 min at 4°C. Protein concentrations were measured using Pierce™ Coomassie (Bradford) Protein Assay Kit (Thermo Fisher Scientific, #23236). Lysates were immunoprecipitated with 15µl pre-washed Pierce™ Anti-HA Magnetic beads (Thermo Fisher Scientific, #88837) at 4°C overnight on a rotor. Beads were washed 4-5 times with Triton X-100 wash buffer (1% Triton X-100, 500 mM NaCl, 50 mM Tris-HCl pH 7.4) and proteins were eluted by incubation at 65°C for 15 min in 2× SDS sample buffer.

For concanavalin A pull-down, N-glycosylated proteins were enriched by incubating cell lysates containing protease inhibitor and 1,10-phenanthroline (Sigma-Aldrich, #131377) with 20µl concanavalin A Sepharose beads (Sigma-Aldrich, # C9017) at 4°C for at least 2 h with rotation. Beads were washed with Triton X-100 wash buffer and proteins were eluted in 2x NuPAGE™ LDS sample buffer (Thermo Fischer Scientific, #NP008) supplemented with 50 mM DTT and 50 % sucrose for 15 min at 65°C and were ran on 4-12 % NuPAGE™ Bis-Tris gels.

### Deglycosylation assay

Cells were lysed in Trition X-100 lysis buffer as described above. Lysates were first denatured with Glycoprotein Denaturing Buffer at 65°C for 15 min and then treated with Endoglycosidase H (Endo H) (New England Biolabs, #P0702S) or Peptide-*N*-Glycosidase F (PNGase F) (New England Biolabs, #P0704S) following the manufacturer’s instructions.

### Quantitative RT-PCR

RNA was isolated from cells using the RNeasy Mini kit (Qiagen, #74104) and reverse transcribed using the SuperScript™ VILO™ cDNA synthesis kit (Thermo Fischer Scientific, #11754050). Resulting cDNA was used for quantitative PCR (qRT-PCR) using the TaqMan™ Gene Expression Master Mix (Applied Biosystems, #4369016). A list of all human TaqMan probes used is provided in Supplementary Table 5. For quantification, the relative quantity of samples was calculated according the comparative △Ct method and normalized to GAPDH. Gene expression was compared to the corresponding wild-type control.

### Immunofluorescence and confocal microscopy

HEK293T cells were plated on 13-mm glass coverslips in 12-well plates and transfected with 100-250 ng of indicated constructs for 48 h prior to fixation. Cells were washed 2 times with PBS and fixed with 4% paraformaldehyde in PBS at room temperature for 20 min. Cells were then washed 3 times with PBS and permeabilised in 0.3% TX-100 in PBS for 20 min. Cells were blocked with 3% BSA in 50 mM Tris pH 7.5 for 30 min after removal of permeabilisation buffer. Cells were incubated overnight with indicated antibodies in blocking buffer at 4°C and then washed 3 times in permeabilisation buffer for (5 min each wash). Coverslips were incubated with corresponding species-specific fluorescently coupled secondary antibodies (Invitrogen) for 30 min. Cells were subsequently washed 4 times with PBS (5 *μ*g/ml DAPI was added in second to last wash), prior to mounting on glass slides with VECTASHIELD® anti-fade mounting medium (Vectorlabs, #H-1000-10). Images were acquired with a laser scanning confocal microscope (Fluoview FV1000; Olympus) with a 60×1.4 NA oil objective and processed using Fiji (Image J).

### Immunohistochemistry

Immunohistochemistry was performed on frozen tissue; sections were air-dried before processed. Tissues were fixed in 4% paraformaldehyde (PFA) at room temperature for 15 min. If PFA fixation was used, samples were permeabilized with 0.1% Triton X-100. Tissues were washed three times with PBS for 5 min each and incubated with 5% goat serum in PBS for 1 h at room temperature to reduce nonspecific binding. Tissues were incubated with primary antibody in 5% goat serum overnight at 4 °C. The following day tissues were washed three times with PBS and incubated with the secondary antibody conjugated with Alexa Fluor (Molecular Probes) in 5% goat serum for 1 h at room temperature. After three washes, sections were incubated for 10 min with DAPI (100 ng/ml). Tissues were mounted onto slides using Immumount (Thermoscientific). Fluorescence was evaluated in one single plane by Zeiss 710 confocal microscopy (Carl Zeiss).

### RNA-Seq

HEK293T cells stably expressing inducible iRhom2-1-374 were treated with 100 ng/ml of doxycycline for 0h, 3h and 6h. Triplicate samples were harvested and RNA was extracted using Direct-zol™ RNA MiniPrep Plus kit (Zymo Research, #R2072) according to the manufacturer’s instructions. PolyA library preparation and RNA sequencing was performed by Novogene (UK) Company Ltd. Paired-end 150 bp sequencing was performed using Illumina NovaSeq 6000 sequencing system.

### RNA-Seq data processing and analysis

RNA-seq data were analysed as previously described (Tellier and Murphy, 2020). Adapters were trimmed with Cutadapt version 1.18 (Martin, 2011) in paired-end mode with the following options: --minimum-length 10 -q 15,10 -j 16 -A GATCGTCGGACTGTAGAACTCTGAAC -a AGATCGGAAGAGCACACGTCTGAACTCCAGTCAC. The remaining rRNA reads were removed by mapping the trimmed reads to the rRNA genes defined in the human ribosomal DNA complete repeating unit (GenBank: U13369.1) with STAR version 2.7.3 (Dobin et al., 2013) and the parameters --runThreadN 16 -- readFilesCommand gunzip -c -k --outReadsUnmapped Fastx --limitBAMsortRAM 20000000000 --outSAMtype BAM SortedByCoordinate. The unmapped reads were mapped to the human GRCh38.p13 reference sequence with STAR version 2.7.3a and the ENCODE parameters: --runThreadN 16 --limitBAMsortRAM 20000000000 --outSAMtype BAM SortedByCoordinate --quantMode GeneCounts --outFilterMultimapNmax 20 --outFilterType BySJout --alignSJoverhangMin 8 --alignSJDBoverhangMin 1 --outFilterMismatchNmax 999 -- alignIntronMin 20 --alignIntronMax 1000000 --alignMatesGapMax 1000000. The number of aligned reads per gene obtained with STAR --quantMode GeneCounts were used to perform the differential expression analysis with DESeq2 version 1.30.1 (Love et al., 2014) and apeglm version 1.18.0 (Zhu et al., 2019). For the iRhom2-1-374 RNA-seq dataset, an adjusted p-value < 0.05 was considered statistically significant. We considered as more likely iRhom2-1-374 target genes those found differentially expressed at both 3 h and 6 h and with either a greater downregulation at 6 h than 3 h or a greater upregulation at 6 h than at 3 h.

Control, non-lesional psoriasis, and lesional psoriasis RNA-seq data were obtained from GSE121212 (Tsoi et al., 2019) and analysed similarly as the iRhom2-1-374 RNA-seq dataset. For the psoriasis dataset, a fold change < -2 or > 2 and an adjusted p-value < 0.05 was considered statistically significant.

Gene ontology (GO) enrichment analysis was performed with the Gene Ontology resource website ((Ashburner et al., 2000); Gene Ontology, 2021). MA plots, heatmaps, and GO plots were produced with GraphPad Prism 9.4.0.

### Statistical analysis and data presentation

All statistical analyses were performed with GraphPad Prism 9.4.0. p-value for RNA-Seq data were analysed by the Wald test, as described in the DESeq2 package (Love et al., 2014). Unless indicated, all experiments were performed 2-3 times. The data are expressed as the mean ± standard error of mean (SEM).

## DATA AVAILABILITY

The data supporting the findings from this study are available within the article files and its supplementary information. RNA-Seq data will be deposited in GEO and accession code will be provided once upload is completed.

## ACKNOWLEDGEMENTS

We thank members of the Freeman lab for their support and advice during the study. We are especially grateful to Sonia Muliyil (University of Oxford) for early help with immunofluorescent staining, Owain Bryant (University of Oxford) for iRhom2 structure illustration predicted by AlphaFold, Shona Murphy (University of Oxford) for support and experimental advice, and Martin Spiess (University of Basel) for generously sharing cavinafungin. This paper was supported by Wellcome Trust Senior Investigator Awards to M.F. (101035/Z/13/Z and 220887/Z/20/Z) and grants awarded to D.P.K. from the Medical Research Council (MR/L010402/1 and MR/S009914/1), Cancer Research UK (C7570/A19107) and the 2016 CHANEL-CERIES research award.

## AUTHOR CONTRIBUTIONS

I.D. designed, performed, and analysed most of the study and wrote the paper. M.T. analysed all RNA-Seq data. A.C., C.M.W. and D.P.K. performed and analysed the immunohistochemistry from patient tissues. A.C. independently validated NLS motifs. C.L. performed and analysed immunofluorescence data. B.Z. performed and analysed data from mouse organs. M.F. analysed and supervised the study, and wrote the paper with I.D. All authors have reviewed the paper.

## CORRESPONDING AUTHORS

Iqbal Dulloo and Matthew Freeman

## FIGURE LEGENDS

**Supplementary Fig. 1.**
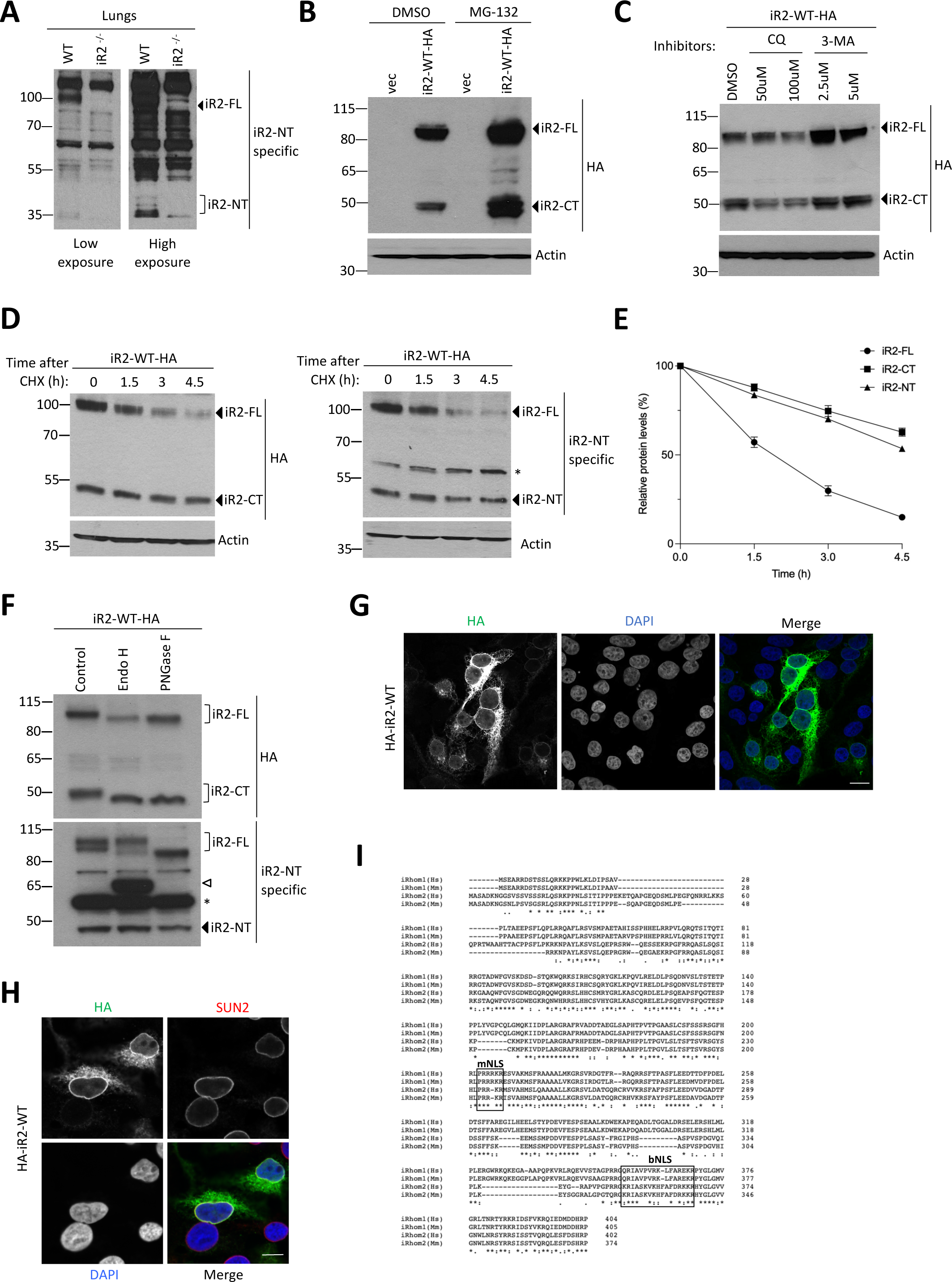
iRhom2 protein fragments are stable and nuclear. **a** Levels of endogenous iRhom2 were analysed by immunoblotting lysates from lung tissues of wild-type and iRhom2 KO mice using iRhom2 N-terminal specific antibody. **b-c** Levels of iRhom2 were analysed by immunoblotting in lysates from HEK293T cells transiently transfected for 36 h with iRhom2-3XHA. Cells were treated for 16h with (**b**) 10 µM MG-132, or (**c**) indicated doses of chloroquine (CQ) or 3-MA. **d** Half-lives of full-length, N-terminal and C-terminal iRhom2 fragments using 100 µg/ml cycloheximide (CHX) were analysed from HEK293T cells transiently transfected for 36 h with iRhom2-3XHA with either HA (left panel) or iRhom2 N-terminal specific antibody (right panel). **e** Quantification of the half-lives of full-length (FL), N-terminal (NT) and C-terminal (CT) iRhom2 fragments (data presented as mean ± SEM; n=3) . **f** A deglycosylation assay was carried out on HEK293T cells transiently transfected for 36 h with iRhom2-3XHA. Lysates were treated with Endo H or PNGase F. Immunoblotting was carried out using either anti-HA or iRhom2 N-terminal specific antibody. * and white arrowhead denote unspecific bands detected with the iRhom2 N-terminal specific antibody. **g-h** Immunofluorescence of N-terminally tagged 3XHA-iRhom2 under a TET-inducible promoter expressed for 18h in cells. Cells were stained for HA (green), DAPI (blue) (**g**) and SUN2 (red) (**h**). Scale bar =10 *μ*m. **i** Alignment of protein sequences of iRhom1 and iRhom2 from *Homo sapiens* (Hs.) and *Mus musculus* (Ms.). Boxed sequences are monopartite (mNLS) or a bipartite (bNLS) nuclear localisation signals, predicted using eukaryotic linear motif (ELM) and PSORT II.

**Supplementary Fig. 2.**
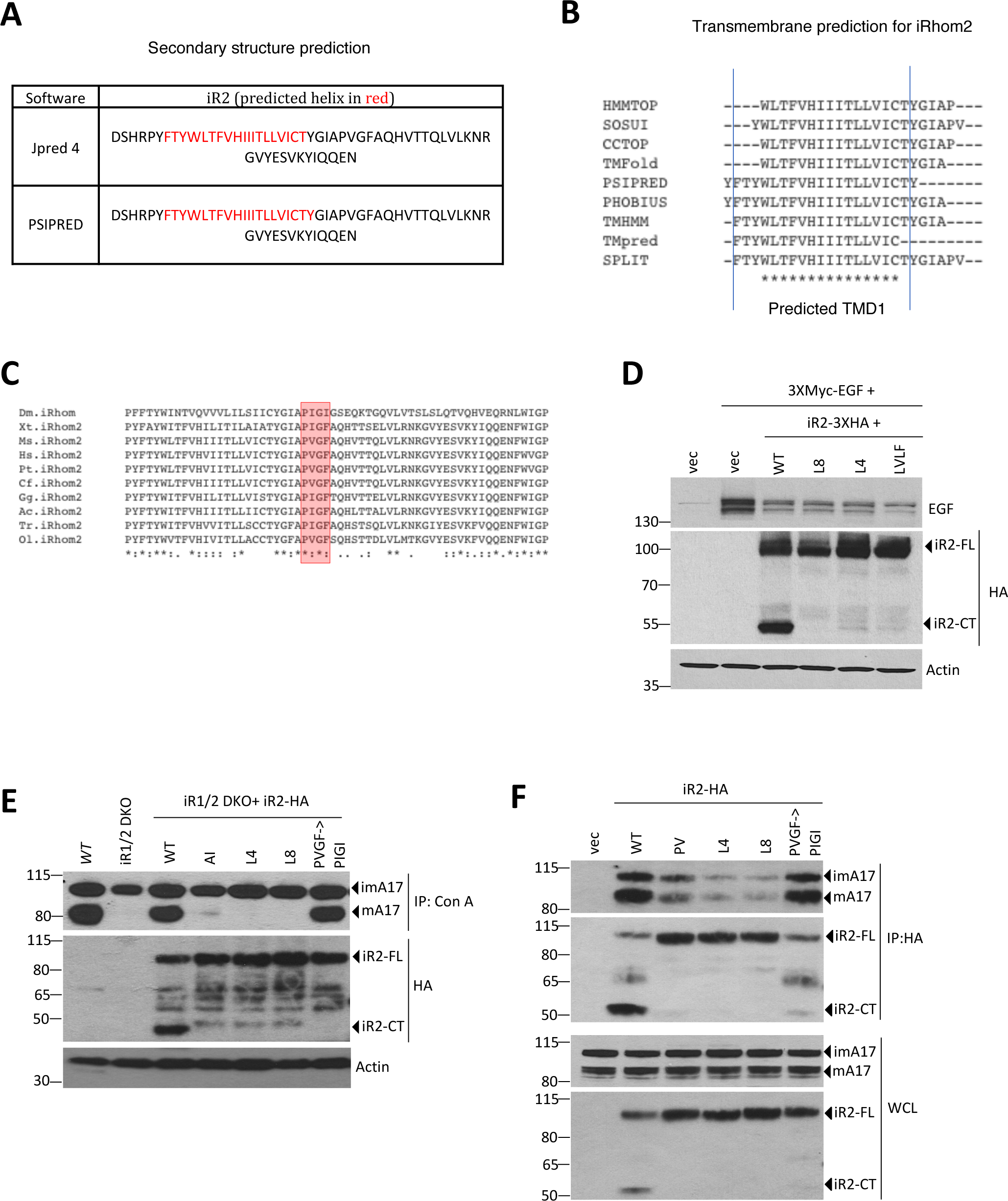
Cleavage of iRhom2 adjacent to TMD1. **a** Secondary structure prediction using JPRED 4 and PSIPRED was carried out on iRhom2 to find the putative boundaries of TM helix 1 (in red). **b** *In silico* analysis of iRhom2 with multiple programs to predict the boundaries of TMD1. **c** Alignment of iRhom2 protein sequences from *Homo sapiens* (Hs.) and *Mus musculus* (Ms), *Drosophila melanogaster* (Dm.), *Xenopus tropicalis* (Xt.), *Pan troglodytes* (Pt.), *Canis familiaris* (Cf.), *Gallus gallus* (Gg.), *Anolis carolinensis* (Ac.), *Takifugu rubripes* (Tr.), *Oryzia latipes* (Ol.). The boxed sequence is the highly conserved four amino acids important for cleavage of iRhoms. **d** Levels of EGF and iRhom2 were analysed by immunoblotting in lysates from HEK293T cells transiently co-transfected for 36 h with iRhom2-3XHA and Myc-EGF. **e** Maturation of ADAM17 using concanavalin A (Con A) pull-down in wild-type, iRhom1/2 DKO, and iRhom1/2 DKO MEFs stably reconstituted with indicated iRhom2 constructs was measured. imA17: immature A17, mA17: mature A17. **f** Levels of iRhom2 and endogenous ADAM17 were determined by immunoblotting in whole cell lysate (WCL) and after immunoprecipitation (IP: HA) from HEK293T cells transfected for 36hrs with the indicated iRhom2 constructs.

**Supplementary Fig. 3.**
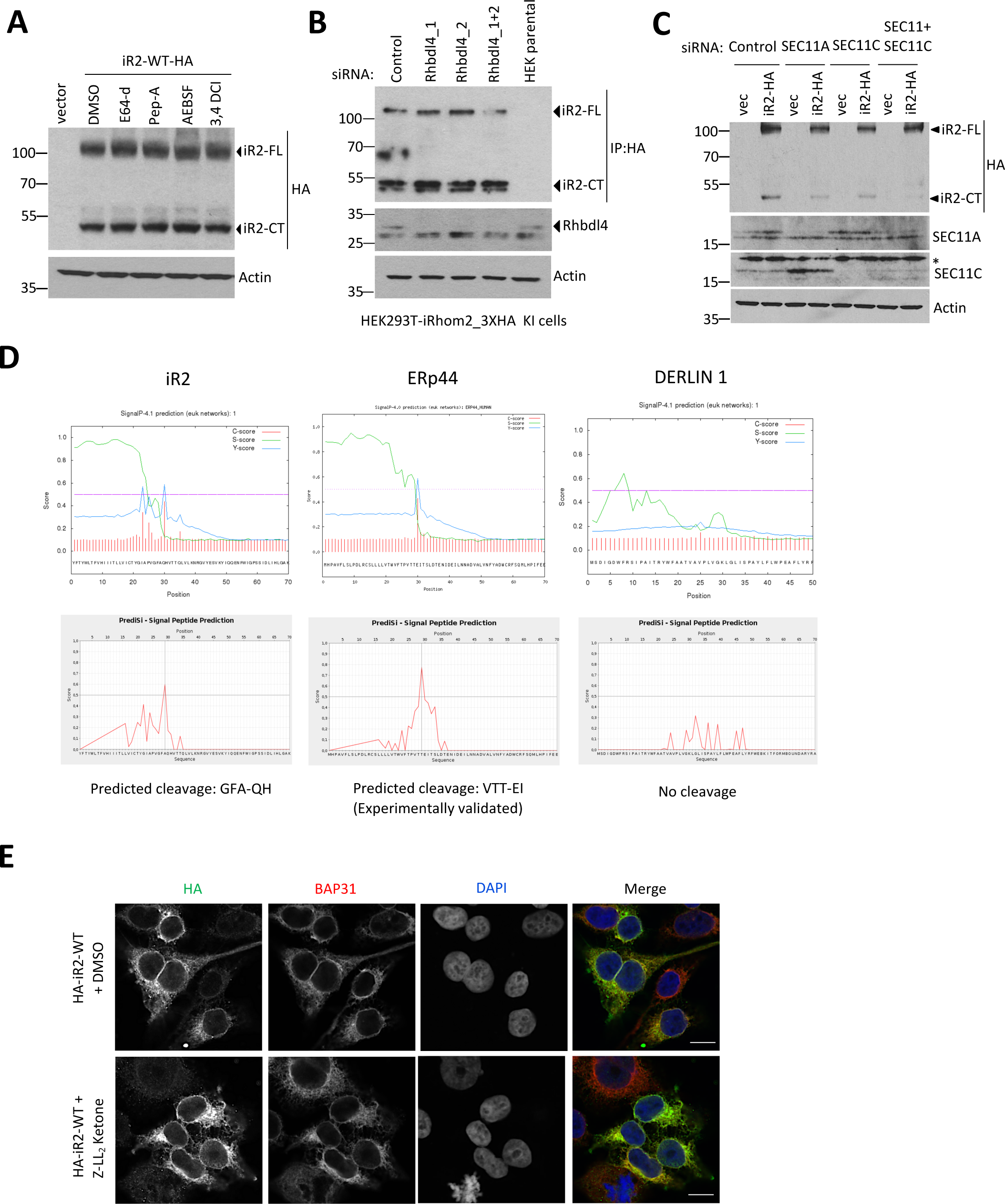
SPC cleaves iRhom2. **a** Levels of iRhom2 were analysed by immunoblotting in lysates from HEK293T cells transiently transfected for 36 h with iRhom2-3XHA in the presence of broad-spectrum protease inhibitors E64-d (20 µM), Pepstatin A (20 µM), AEBSF (200 µM), 3,4 DCI (20 µM) for 16hrs prior to harvest. **b-c** Levels of endogenous iRhom2 (**b**) or transfected iRhom2-3XHA (**c**) were detected by immunoblotting in lysates from HEK293T cells transfected for 72 h with control siRNA or siRNA against either RHBLD4 (**b**) or the catalytic subunits of SPC (**c**). **d** Predicted site of signal peptidase cleavage of iRhom2 by SignalP 4.1 and PrediSi, compared with to experimentally validated ERp44 as a positive control and the rhomboid pseudoprotease DERLIN1 as a negative control. **e** Immunofluorescence of N-terminally tagged 3XHA-iRhom2 under a TET-inducible promoter expressed for 18h in the presence of SPP inhibitor Z-LL_2_ ketone (50 µM). Cells were stained for HA (green), BAP31 (red) and DAPI (blue). Scale bar =10 *μ*m.

**Supplementary Fig 4.**
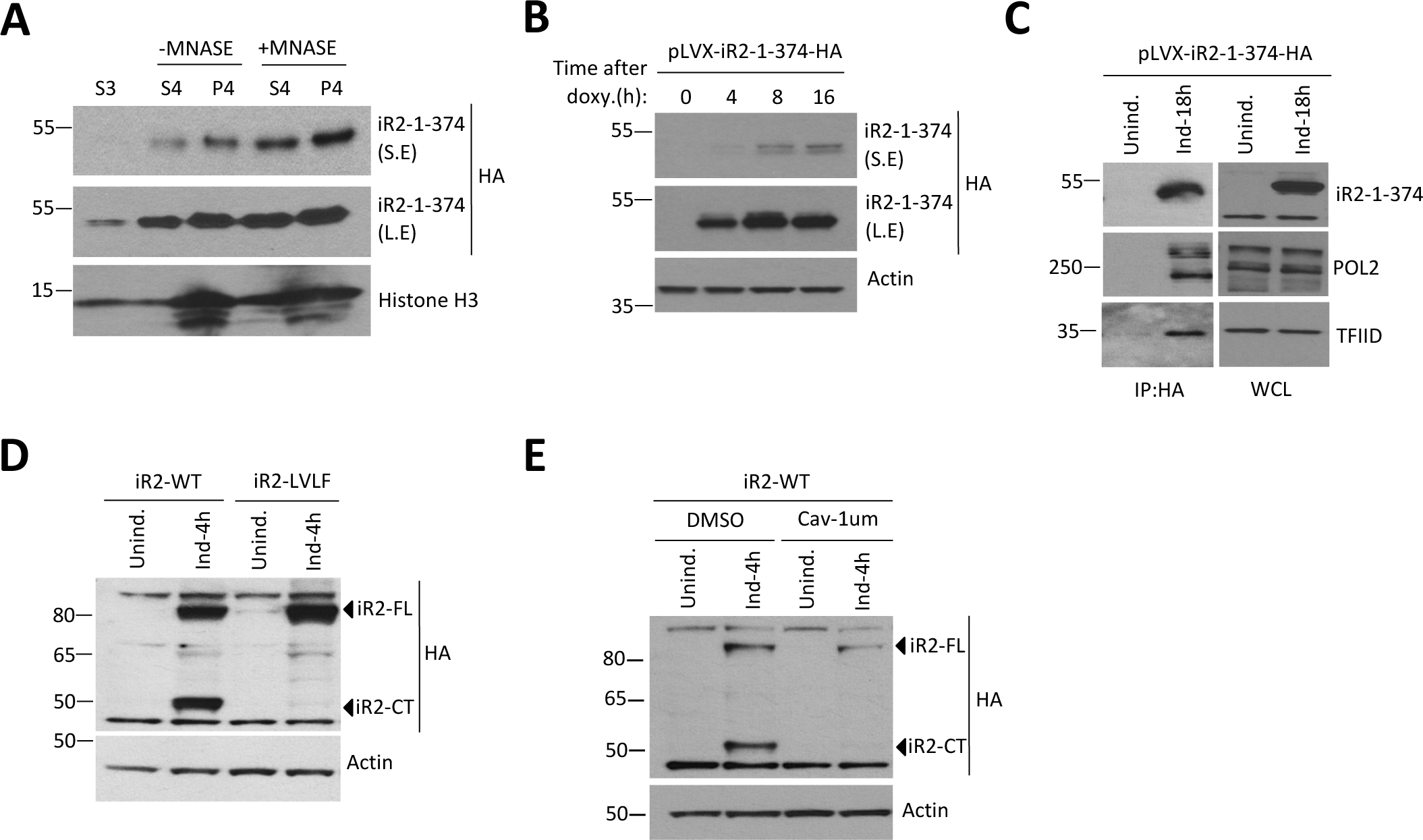
Gene expression regulation by nuclear iRhom2. **a** Levels of nuclear iRhom2 were determined by subcellular fractionation in HEK293T cells stably expressing iR2-1-374. S3 denotes soluble nuclear fraction, S4 denotes chromatin-bound fraction with or without MNASE treatment, and P4 denotes cytoskeletal proteins + remaining chromatin-bound proteins. Histone H3 is a positive control for chromatin-bound proteins **b** Levels of iR2-1-374 were analysed by immunoblotting in HEK293T cells expressing iR2-1-374 under a TET-inducible promoter for indicated periods. **c** Levels of POL2 and TFIID were analysed by immunoblotting in whole cell lysate (WCL) and immunoprecipitation (IP: HA) from HEK293T cells expressing iR2-1-374 under a TET-inducible promoter for 18 h. **d-e** Immunoblot analysis of HEK293T cells expressing for 4 h either inducible wild-type (iR2-WT) or uncleavable iRhom2 (iR2-LVLF) (**d**) or inducible wild-type iRhom2 in the presence of cavinafungin (Cav-1 µM) (**e**). S.E and L.E. denote short and long exposure respectively.

**Supplementary Fig 5.**
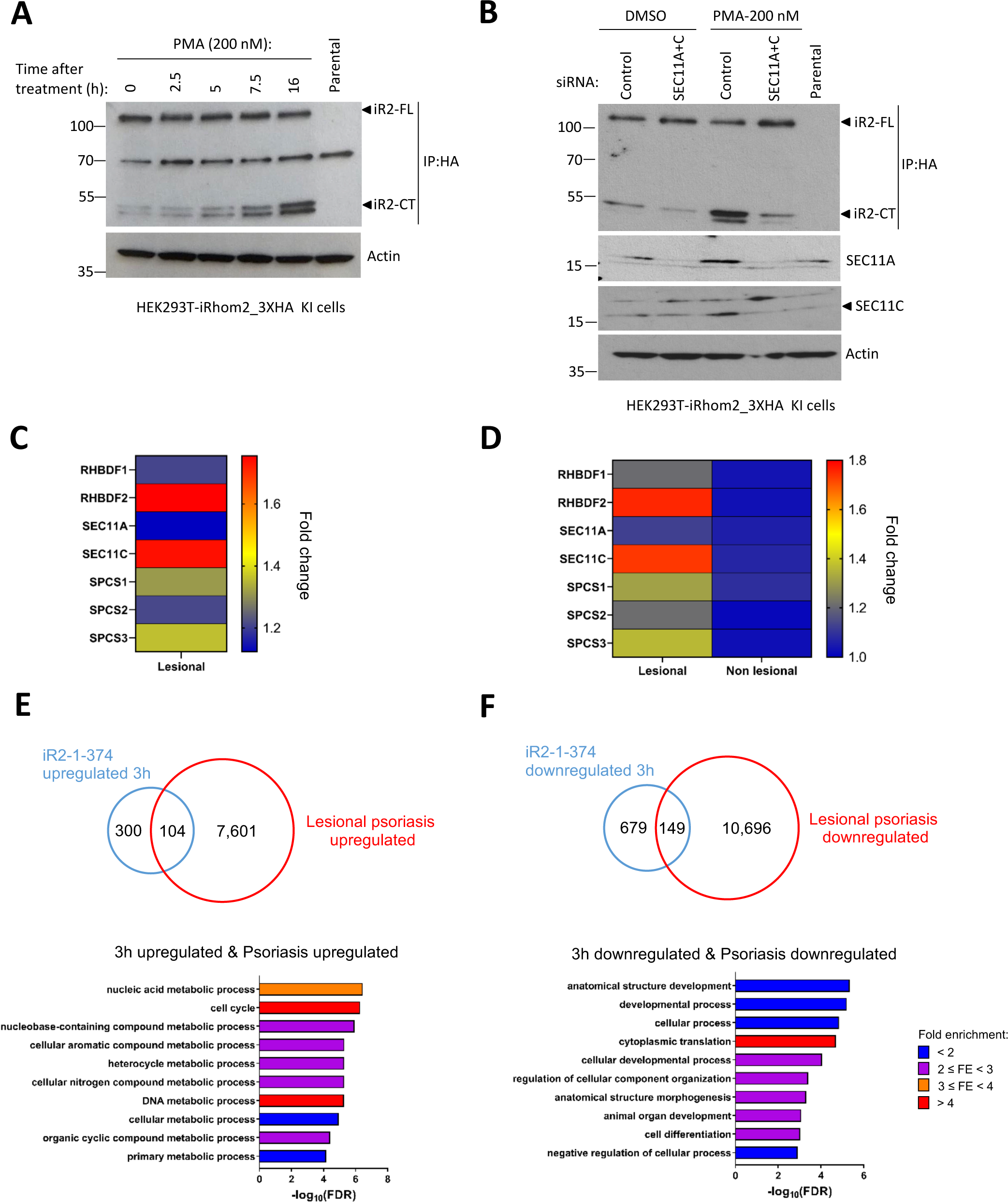
Expression and regulation of nuclear iRhom2 in disease skin models. **a-b** Levels of endogenous iRhom2 were analysed by immunoblotting in lysates from cells treated with 200 nM PMA for the indicated time intervals (**a**) or after transfection for 72 h with either control siRNA or siRNA against SEC11A and SEC11C (**b**). **c-d** Fold change of gene expression from a published RNA-Seq dataset of iRhom1 (RHBDF1), iRhom2 (RHBDF2) and all subunits of SPC (SEC11A, SEC11B, SPCS1, SPCS2, SPCS3) in lesional psoriasis compared to control (**c**) or lesional psoriasis compared non-lesional psoriasis (**d**). **e-f** Venn diagrams showing overlap between differentially expressed genes found in a published RNA-Seq dataset of lesional psoriasis and upregulated (**e**) or downregulated (**f**) at 3 h induction of iR2-1-374 (top). Bar graphs showing summary of the GO enrichment analysis on the set of common upregulated (n=104) or downregulated (n=149) genes between lesional psoriasis and iR2-1-374. FDR: False discovery rate. Different colours denote range of fold enrichment (FE).

## REFERENCES

1. Adrain, C., and Freeman, M. (2012). New lives for old: evolution of pseudoenzyme function illustrated by iRhoms. Nat Rev Mol Cell Biol 13, 489–498.

2. Adrain, C., Zettl, M., Christova, Y., Taylor, N., and Freeman, M. (2012). Tumor necrosis factor signaling requires iRhom2 to promote trafficking and activation of TACE. Science 335, 225–228.

3. Al-Salihi, M.A., and Lang, P.A. (2020). iRhom2: An Emerging Adaptor Regulating Immunity and Disease. Int J Mol Sci 21.

4. Alzahrani, N., Wu, M.J., Shanmugam, S., and Yi, M. (2020). Delayed by Design: Role of Suboptimal Signal Peptidase Processing of Viral Structural Protein Precursors in Flaviviridae Virus Assembly. Viruses 12.

5. Ashburner, M., Ball, C.A., Blake, J.A., Botstein, D., Butler, H., Cherry, J.M., Davis, A.P., Dolinski, K., Dwight, S.S., Eppig, J.T., et al. (2000). Gene ontology: tool for the unification of biology. The Gene Ontology Consortium. Nat Genet 25, 25–29.

6. Blaydon, D.C., Etheridge, S.L., Risk, J.M., Hennies, H.C., Gay, L.J., Carroll, R., Plagnol, V., McRonald, F.E., Stevens, H.P., Spurr, N.K., et al. (2012). RHBDF2 mutations are associated with tylosis, a familial esophageal cancer syndrome. Am J Hum Genet 90, 340–346.

7. Brooke, M.A., Etheridge, S.L., Kaplan, N., Simpson, C., O’Toole, E.A., Ishida-Yamamoto, A., Marches, O., Getsios, S., and Kelsell, D.P. (2014). iRHOM2-dependent regulation of ADAM17 in cutaneous disease and epidermal barrier function. Hum Mol Genet 23, 4064–4076.

8. Cai, M., Huang, Y., Ghirlando, R., Wilson, K.L., Craigie, R., and Clore, G.M. (2001). Solution structure of the constant region of nuclear envelope protein LAP2 reveals two LEM-domain structures: one binds BAF and the other binds DNA. EMBO J 20, 4399–4407.

9. Caputo, S., Couprie, J., Duband-Goulet, I., Konde, E., Lin, F., Braud, S., Gondry, M., Gilquin, B., Worman, H.J., and Zinn-Justin, S. (2006). The carboxyl-terminal nucleoplasmic region of MAN1 exhibits a DNA binding winged helix domain. J Biol Chem 281, 18208–18215.

10. Chang, S.N., Dey, D.K., Oh, S.T., Kong, W.H., Cho, K.H., Al-Olayan, E.M., Hwang, B.S., Kang, S.C., and Park, J.G. (2020). Phorbol 12-Myristate 13-Acetate Induced Toxicity Study and the Role of Tangeretin in Abrogating HIF-1alpha-NF-kappaB Crosstalk In Vitro and In Vivo. Int J Mol Sci 21.

11. Choo, K.H., and Ranganathan, S. (2008). Flanking signal and mature peptide residues influence signal peptide cleavage. BMC Bioinformatics 9 *Suppl 12*, S15.

12. Christova, Y., Adrain, C., Bambrough, P., Ibrahim, A., and Freeman, M. (2013). Mammalian iRhoms have distinct physiological functions including an essential role in TACE regulation. EMBO Rep 14, 884–890.

13. Czapiewski, R., Robson, M.I., and Schirmer, E.C. (2016). Anchoring a Leviathan: How the Nuclear Membrane Tethers the Genome. Frontiers in genetics 7, 82.

14. De Strooper, B., Annaert, W., Cupers, P., Saftig, P., Craessaerts, K., Mumm, J.S., Schroeter, E.H., Schrijvers, V., Wolfe, M.S., Ray, W.J., et al. (1999). A presenilin-1-dependent gamma-secretase-like protease mediates release of Notch intracellular domain. Nature 398, 518–522.

15. Dobin, A., Davis, C.A., Schlesinger, F., Drenkow, J., Zaleski, C., Jha, S., Batut, P., Chaisson, M., and Gingeras, T.R. (2013). STAR: ultrafast universal RNA-seq aligner. Bioinformatics 29, 15–21.

16. Dulloo, I., Atakpa-Adaji, P., Yeh, Y.C., Levet, C., Muliyil, S., Lu, F., Taylor, C.W., and Freeman, M. (2022). iRhom pseudoproteases regulate ER stress-induced cell death through IP(3) receptors and BCL-2. Nat Commun 13, 1257.

17. Dulloo, I., Muliyil, S., and Freeman, M. (2019). The molecular, cellular and pathophysiological roles of iRhom pseudoproteases. Open biology 9, 190003.

18. Estoppey, D., Lee, C.M., Janoschke, M., Lee, B.H., Wan, K.F., Dong, H., Mathys, P., Filipuzzi, I., Schuhmann, T., Riedl, R., et al. (2017). The Natural Product Cavinafungin Selectively Interferes with Zika and Dengue Virus Replication by Inhibition of the Host Signal Peptidase. Cell Rep 19, 451–460.

19. Fleig, L., Bergbold, N., Sahasrabudhe, P., Geiger, B., Kaltak, L., and Lemberg, M.K. (2012). Ubiquitin-dependent intramembrane rhomboid protease promotes ERAD of membrane proteins. Mol Cell 47, 558–569.

20. Freeman, M. (2014). The rhomboid-like superfamily: molecular mechanisms and biological roles. Annu Rev Cell Dev Biol 30, 235–254.

21. Grieve, A.G., Xu, H., Kunzel, U., Bambrough, P., Sieber, B., and Freeman, M. (2017). Phosphorylation of iRhom2 at the plasma membrane controls mammalian TACE-dependent inflammatory and growth factor signalling. Elife 6.

22. Griffiths, C.E., and Barker, J.N. (2007). Pathogenesis and clinical features of psoriasis. Lancet 370, 263–271.

23. Hegde, R.S., and Bernstein, H.D. (2006). The surprising complexity of signal sequences. Trends Biochem Sci 31, 563–571.

24. Hiller, K., Grote, A., Scheer, M., Munch, R., and Jahn, D. (2004). PrediSi: prediction of signal peptides and their cleavage positions. Nucleic Acids Res 32, W375–379.

25. Hosur, V., Johnson, K.R., Burzenski, L.M., Stearns, T.M., Maser, R.S., and Shultz, L.D. (2014). Rhbdf2 mutations increase its protein stability and drive EGFR hyperactivation through enhanced secretion of amphiregulin. Proc Natl Acad Sci U S A 111, E2200–2209.

26. Hosur, V., Low, B.E., Shultz, L.D., and Wiles, M.V. (2017a). Genetic deletion of amphiregulin restores the normal skin phenotype in a mouse model of the human skin disease tylosis. Biol Open 6, 1174–1179.

27. Hosur, V., Lyons, B.L., Burzenski, L.M., and Shultz, L.D. (2017b). Tissue-specific role of RHBDF2 in cutaneous wound healing and hyperproliferative skin disease. BMC Res Notes 10, 573.

28. Jackson, R.C., and Blobel, G. (1977). Post-translational cleavage of presecretory proteins with an extract of rough microsomes from dog pancreas containing signal peptidase activity. Proc Natl Acad Sci U S A 74, 5598–5602.

29. Kilic, A., Klose, S., Dobberstein, B., Knust, E., and Kapp, K. (2010). The Drosophila Crumbs signal peptide is unusually long and is a substrate for signal peptide peptidase. Eur J Cell Biol 89, 449–461.

30. Kunzel, U., Grieve, A.G., Meng, Y., Sieber, B., Cowley, S.A., and Freeman, M. (2018). FRMD8 promotes inflammatory and growth factor signalling by stabilising the iRhom/ADAM17 sheddase complex. Elife 7.

31. Lemberg, M.K., and Adrain, C. (2016). Inactive rhomboid proteins: New mechanisms with implications in health and disease. Semin Cell Dev Biol 60, 29–37.

32. Lemberg, M.K., and Freeman, M. (2007). Functional and evolutionary implications of enhanced genomic analysis of rhomboid intramembrane proteases. Genome Res 17, 1634–1646.

33. Liaci, A.M., Steigenberger, B., Telles de Souza, P.C., Tamara, S., Grollers-Mulderij, M., Ogrissek, P., Marrink, S.J., Scheltema, R.A., and Forster, F. (2021). Structure of the human signal peptidase complex reveals the determinants for signal peptide cleavage. Mol Cell 81, 3934–3948 e3911.

34. Lin, F., Morrison, J.M., Wu, W., and Worman, H.J. (2005). MAN1, an integral protein of the inner nuclear membrane, binds Smad2 and Smad3 and antagonizes transforming growth factor-beta signaling. Hum Mol Genet 14, 437–445.

35. Love, M.I., Huber, W., and Anders, S. (2014). Moderated estimation of fold change and dispersion for RNA-seq data with DESeq2. Genome Biol 15, 550.

36. Lowes, M.A., Suarez-Farinas, M., and Krueger, J.G. (2014). Immunology of psoriasis. Annu Rev Immunol 32, 227–255.

37. Luo, W.W., Li, S., Li, C., Lian, H., Yang, Q., Zhong, B., and Shu, H.B. (2016). iRhom2 is essential for innate immunity to DNA viruses by mediating trafficking and stability of the adaptor STING. Nat Immunol 17, 1057–1066.

38. Maney, S.K., McIlwain, D.R., Polz, R., Pandyra, A.A., Sundaram, B., Wolff, D., Ohishi, K., Maretzky, T., Brooke, M.A., Evers, A., et al. (2015). Deletions in the cytoplasmic domain of iRhom1 and iRhom2 promote shedding of the TNF receptor by the protease ADAM17. Sci Signal 8, ra109.

39. Martin, M. (2011). Cutadapt removes adapter sequences from high-throughput sequencing reads. 2011 *17*, 3.

40. Maruthappu, T., Chikh, A., Fell, B., Delaney, P.J., Brooke, M.A., Levet, C., Moncada-Pazos, A., Ishida-Yamamoto, A., Blaydon, D., Waseem, A., et al. (2017). Rhomboid family member 2 regulates cytoskeletal stress-associated Keratin 16. Nat Commun 8, 14174.

41. Massague, J., Seoane, J., and Wotton, D. (2005). Smad transcription factors. Genes Dev 19, 2783–2810.

42. McIlwain, D.R., Lang, P.A., Maretzky, T., Hamada, K., Ohishi, K., Maney, S.K., Berger, T., Murthy, A., Duncan, G., Xu, H.C., et al. (2012). iRhom2 regulation of TACE controls TNF-mediated protection against Listeria and responses to LPS. Science 335, 229–232.

43. Meinema, A.C., Laba, J.K., Hapsari, R.A., Otten, R., Mulder, F.A., Kralt, A., van den Bogaart, G., Lusk, C.P., Poolman, B., and Veenhoff, L.M. (2011). Long unfolded linkers facilitate membrane protein import through the nuclear pore complex. Science 333, 90–93.

44. Mentrup, T., Fluhrer, R., and Schroder, B. (2017). Latest emerging functions of SPP/SPPL intramembrane proteases. Eur J Cell Biol 96, 372–382.

45. Mudumbi, K.C., Czapiewski, R., Ruba, A., Junod, S.L., Li, Y., Luo, W., Ngo, C., Ospina, V., Schirmer, E.C., and Yang, W. (2020). Nucleoplasmic signals promote directed transmembrane protein import simultaneously via multiple channels of nuclear pores. Nat Commun 11, 2184.

46. Nakagawa, T., Guichard, A., Castro, C.P., Xiao, Y., Rizen, M., Zhang, H.Z., Hu, D., Bang, A., Helms, J., Bier, E., et al. (2005). Characterization of a human rhomboid homolog, p100hRho/RHBDF1, which interacts with TGF-alpha family ligands. Dev Dyn 233, 1315–1331.

47. Nestle, F.O., Kaplan, D.H., and Barker, J. (2009). Psoriasis. N Engl J Med 361, 496–509.

48. Oikonomidi, I., Burbridge, E., Cavadas, M., Sullivan, G., Collis, B., Naegele, H., Clancy, D., Brezinova, J., Hu, T., Bileck, A., et al. (2018). iTAP, a novel iRhom interactor, controls TNF secretion by policing the stability of iRhom/TACE. Elife 7.

49. Owji, H., Nezafat, N., Negahdaripour, M., Hajiebrahimi, A., and Ghasemi, Y. (2018). A comprehensive review of signal peptides: Structure, roles, and applications. Eur J Cell Biol 97, 422–441.

50. Pan, D., Estevez-Salmeron, L.D., Stroschein, S.L., Zhu, X., He, J., Zhou, S., and Luo, K. (2005). The integral inner nuclear membrane protein MAN1 physically interacts with the R-Smad proteins to repress signaling by the transforming growth factor-beta superfamily of cytokines. J Biol Chem 280, 15992–16001.

51. Paschkowsky, S., Hamze, M., Oestereich, F., and Munter, L.M. (2016). Alternative Processing of the Amyloid Precursor Protein Family by Rhomboid Protease RHBDL4. J Biol Chem 291, 21903–21912.

52. Pasquali, L., Srivastava, A., Meisgen, F., Das Mahapatra, K., Xia, P., Xu Landen, N., Pivarcsi, A., and Sonkoly, E. (2019). The Keratinocyte Transcriptome in Psoriasis: Pathways Related to Immune Responses, Cell Cycle and Keratinization. Acta dermato-venereologica 99, 196–205.

53. Petersen, T.N., Brunak, S., von Heijne, G., and Nielsen, H. (2011). SignalP 4.0: discriminating signal peptides from transmembrane regions. Nat Methods 8, 785–786.

54. Pla-Prats, C., and Thoma, N.H. (2022). Quality control of protein complex assembly by the ubiquitin-proteasome system. Trends Cell Biol 32, 696–706.

55. Ribeiro, A.J.M., Das, S., Dawson, N., Zaru, R., Orchard, S., Thornton, J.M., Orengo, C., Zeqiraj, E., Murphy, J.M., and Eyers, P.A. (2019). Emerging concepts in pseudoenzyme classification, evolution, and signaling. Sci Signal 12.

56. Roeder, R.G. (2019). 50+ years of eukaryotic transcription: an expanding universe of factors and mechanisms. Nat Struct Mol Biol 26, 783–791.

57. Sakai, J., Duncan, E.A., Rawson, R.B., Hua, X., Brown, M.S., and Goldstein, J.L. (1996). Sterol-regulated release of SREBP-2 from cell membranes requires two sequential cleavages, one within a transmembrane segment. Cell 85, 1037–1046.

58. Shelness, G.S., and Blobel, G. (1990). Two subunits of the canine signal peptidase complex are homologous to yeast SEC11 protein. J Biol Chem 265, 9512–9519.

59. Siggs, O.M., Grieve, A., Xu, H., Bambrough, P., Christova, Y., and Freeman, M. (2014). Genetic interaction implicates iRhom2 in the regulation of EGF receptor signalling in mice. Biol Open 3, 1151–1157.

60. Snapp, E.L., McCaul, N., Quandte, M., Cabartova, Z., Bontjer, I., Kallgren, C., Nilsson, I., Land, A., von Heijne, G., Sanders, R.W., et al. (2017). Structure and topology around the cleavage site regulate post-translational cleavage of the HIV-1 gp160 signal peptide. Elife 6.

61. Subramanian, A., Tamayo, P., Mootha, V.K., Mukherjee, S., Ebert, B.L., Gillette, M.A., Paulovich, A., Pomeroy, S.L., Golub, T.R., Lander, E.S., et al. (2005). Gene set enrichment analysis: a knowledge-based approach for interpreting genome-wide expression profiles. Proc Natl Acad Sci U S A 102, 15545–15550.

62. Tellier, M., and Murphy, S. (2020). Incomplete removal of ribosomal RNA can affect chromatin RNA-seq data analysis. Transcription 11, 230–235.

63. Tsoi, L.C., Rodriguez, E., Degenhardt, F., Baurecht, H., Wehkamp, U., Volks, N., Szymczak, S., Swindell, W.R., Sarkar, M.K., Raja, K., et al. (2019). Atopic Dermatitis Is an IL-13-Dominant Disease with Greater Molecular Heterogeneity Compared to Psoriasis. J Invest Dermatol 139, 1480–1489.

64. Ulbrecht, M., Martinozzi, S., Grzeschik, M., Hengel, H., Ellwart, J.W., Pla, M., and Weiss, E.H. (2000). Cutting edge: the human cytomegalovirus UL40 gene product contains a ligand for HLA-E and prevents NK cell-mediated lysis. J Immunol 164, 5019–5022.

65. von Heijne, G. (1983). Patterns of amino acids near signal-sequence cleavage sites. Eur J Biochem 133, 17–21.

66. von Heijne, G. (1984). Analysis of the distribution of charged residues in the N-terminal region of signal sequences: implications for protein export in prokaryotic and eukaryotic cells. EMBO J 3, 2315–2318.

67. von Messling, V., and Cattaneo, R. (2002). Amino-terminal precursor sequence modulates canine distemper virus fusion protein function. J Virol 76, 4172–4180.

68. Weihofen, A., Binns, K., Lemberg, M.K., Ashman, K., and Martoglio, B. (2002). Identification of signal peptide peptidase, a presenilin-type aspartic protease. Science 296, 2215–2218.

69. Wysocka, J., Reilly, P.T., and Herr, W. (2001). Loss of HCF-1-chromatin association precedes temperature-induced growth arrest of tsBN67 cells. Mol Cell Biol 21, 3820–3829.

70. Xie, S., Chen, Z., Wang, Q., Song, X., and Zhang, L. (2014). Comparisons of gene expression in normal, lesional, and non-lesional psoriatic skin using DNA microarray techniques. Int J Dermatol 53, 1213–1220.

71. Ye, J., Rawson, R.B., Komuro, R., Chen, X., Dave, U.P., Prywes, R., Brown, M.S., and Goldstein, J.L. (2000). ER stress induces cleavage of membrane-bound ATF6 by the same proteases that process SREBPs. Mol Cell 6, 1355–1364.

72. Zettl, M., Adrain, C., Strisovsky, K., Lastun, V., and Freeman, M. (2011). Rhomboid family pseudoproteases use the ER quality control machinery to regulate intercellular signaling. Cell 145, 79–91.

73. Zhang, G., Liu, X., Wang, C., Qu, L., Deng, J., Wang, H., and Qin, Z. (2013). Resolution of PMA-induced skin inflammation involves interaction of IFN-gamma and ALOX15. Mediators Inflamm 2013, 930124.

74. Zhu, A., Ibrahim, J.G., and Love, M.I. (2019). Heavy-tailed prior distributions for sequence count data: removing the noise and preserving large differences. Bioinformatics 35, 2084–2092.

75. Zunke, F., and Rose-John, S. (2017). The shedding protease ADAM17: Physiology and pathophysiology. Biochimica et biophysica acta Molecular cell research 1864, 2059–2070.

